# The integrated role of resource memory and scent-based territoriality in the emergence of home-ranges

**DOI:** 10.1101/2021.05.07.443202

**Authors:** Meryl Theng, Thomas A. A. Prowse, Steven Delean, Phillip Cassey, Chloe Bracis

## Abstract

Despite decades of animal movement research, an integrated understanding of the processes underlying the emergence of home-ranges remains inadequate. We explored the effects of integrating two key processes of home-ranging arising from individual movement decisions: (i) memory-based resource use; and (ii) scent-based territoriality. Both mechanisms together led to the formation of exclusive, resource- and population density-dependent home-ranges, which were responsive to perturbations in the conspecific environment (i.e., removing individuals). Home-range patterns (size, size skewness and overlap) were primarily influenced by memory and scent decay rate parameters, demonstrating that home-ranging is ultimately a balance between an animal’s inherent exploratory tendency and its desire to avoid conspecifics. Model application to a population of feral cats demonstrated that general space use patterns could be approximated through simulation, and replication of finer-scale space use patterns is plausible with further model development. Our modelling framework provides a foundation for sophisticated theoretical models of space use in interacting animals.

## 1 Introduction

Many animal species constrain their movement to specific home-ranges (HR), which emerge from behaviours inherently linked to survival and reproduction (Burt 1943). The emergent HRs result from movement decisions of individual animals, which are driven by dynamic processes including their need to access resources while avoiding costly interactions with conspecifics and predators (Borger *et al*. 2008; Nathan *et al*. 2008). Despite decades of extensive research, it is only recently that models of HR emergence have been developed (Ranc *et al*. 2020). Two key processes are proposed to play important roles in HR formation in territorial animals: (1) optimising resource acquisition by referencing a cognitive memory (i.e., resource memory); and (2) minimising resource competition through defensive cues (i.e., territoriality) (Borger *et al*. 2008; Powell & Mitchell 2012; Spencer 2012; Fagan *et al*. 2013). While there have been successes in modelling these underlying mechanisms separately, integration of the two to form a general predictive theory of HR emergence has remained a key challenge (Potts & Lewis 2014).

A fundamental characteristic of animal HRs is the regular revisitation to locations such as foraging areas, dens, watering holes and movement corridors (a.k.a. ‘site fidelity’). Animal memory provides a plausible biological explanation of this phenomenon and recent empirical evidence supports this hypothesis (Bracis & Mueller 2017; Merkle *et al*. 2017; Ranc *et al*. 2021). While quantifying memory is particularly challenging, theoretical analyses have demonstrated that memory-based foraging processes can produce emergent home ranges and more efficient resource use (Van Moorter *et al*. 2009; Bracis *et al*. 2015; Riotte-Lambert *et al*. 2015). Modelling memory mechanisms essentially captures the underlying process behind formation of HR boundaries, and spatio-temporal patterns of site use and fidelity within a HR. It allows replication of the dynamic nature of HRs as a response to a changing environment (e.g., Potts *et al*. 2013; Bateman *et al*. 2015), which is a key advance from non-mechanistic models that have commonly assigned localising centres or HR boundaries to achieve stable, but unrealistically static HRs (Borger *et al*. 2008).

Competitive interactions can drive spatial segregation of HRs, particularly in territorial animals that maintain and defend exclusive territories against conspecifics. In classical mechanistic models of animal movement, territoriality is modelled as scent-mediated conspecific avoidance (Giuggioli *et al*. 2013; Potts & Lewis 2014), which has been demonstrated as a significant underlying driver of observed variation in individual HRs and changes in HR patterns following population change in territorial carnivores (Lewis & Murray 1993; Moorcroft *et al*. 2006; Bateman *et al*. 2015). While these models have led to realistic patterns of HR formation, most have imposed a redirect-to-centre response to scent to stabilise otherwise unconstrained expansion of HRs caused by diffusive movement (Borger *et al*. 2008; Potts & Lewis 2014). This non-mechanistic component is not suitable for modelling HRs of animals that are not central place foragers or denning animals, nor does it allow the emergence of dynamic localising behaviours as a response to changing environments (e.g., HR shifts following resource depletion or conspecific removal). Moreover, the redirect-to-centre response does not explain the underlying localising movement behaviours in the absence of conspecifics (e.g., in sparsely populated habitats), which can result from memory processes.

Though critical insight has been gained from modelling resource memory and territoriality separately, the integration of these two important aspects of HR formation is unexplored. Each component essentially provides a mechanistic explanation for what the other lacks: resource memory is an attractive driver for individuals to preferentially acquire resources from previously visited sites, while territoriality is a repulsive driver which causes individuals to establish exclusive HRs in a multi-individual context (Potts & Lewis 2014). Here, we advance toward a more holistic understanding of HR formation, by developing what is, to our knowledge, the first theoretical model integrating resource memory and territorial processes to simulate realistic patterns of space use by territorial animals. We extend a memory-based model (Bracis *et al*. 2015) to include territoriality. To explore and illustrate the effects of integrating these two components, we investigate how the interplay of resource memory and territoriality influences: (1) the emergence of individual HRs; (2) the relationship between HR size, population density and resource availability; and (3) the response of animal HR to changes in the conspecific environment (i.e., removing individuals). Next, we conduct a sensitivity analysis on the movement model to assess parameter importance for HR patterns (size, size skewness and overlap) and other metrics of movement (step-length, step-length distribution skewness, daily distance) to guide the calibration of model parameters using telemetry data.

## 2 Methods

We first describe the conceptual movement model (**Section 2.1**; summarised in **Fig. 1**), and then detail the parameterisation process used for simulations in the following **Sections 2.2** and **2.3**. All parameter values used in simulations are collated in **Table 1**.

**Figure 1.**
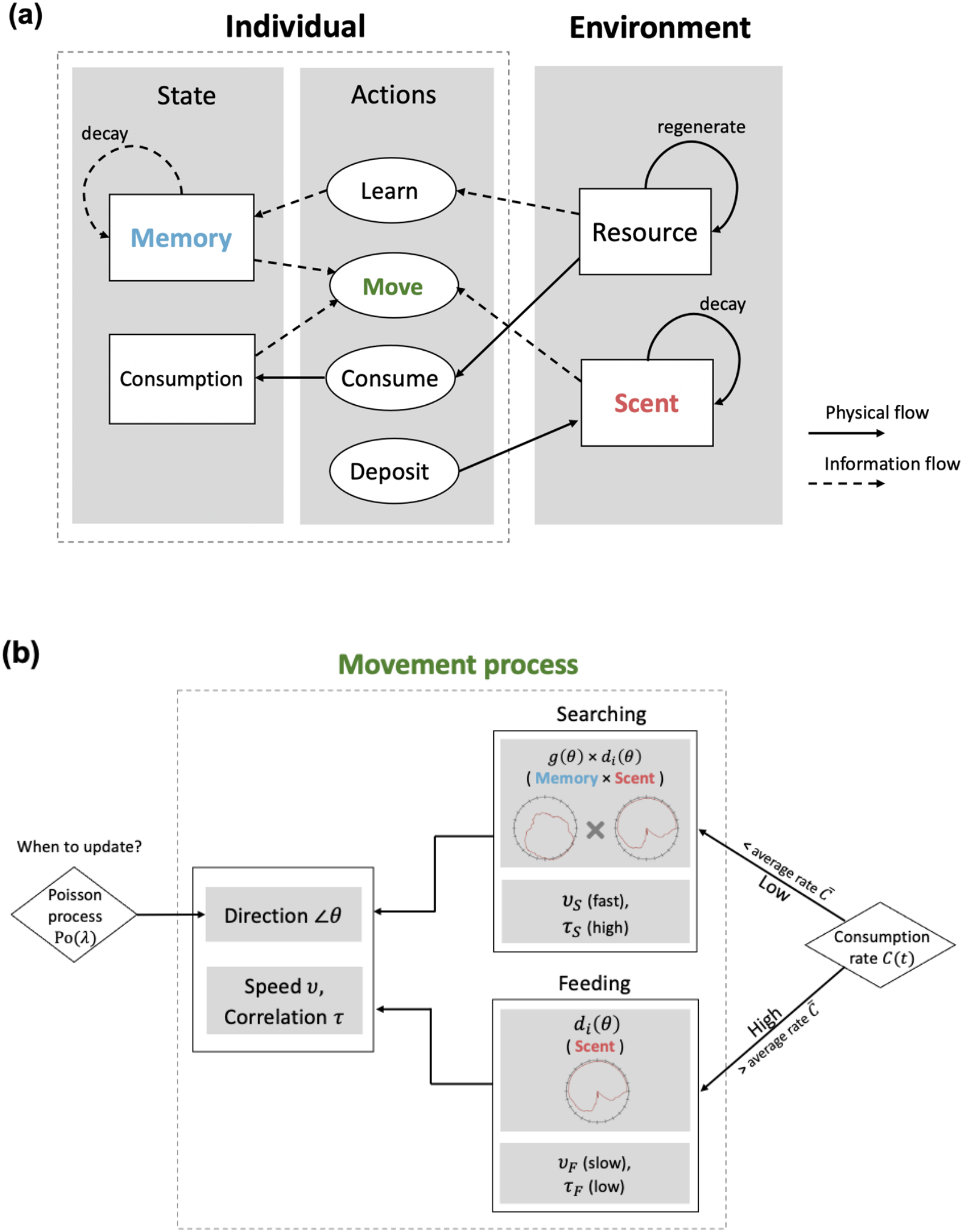
A conceptual representation of the movement model. a) An overview of the individual-environment processes. The individual interacts with the external environment through a series of actions, which serve to update its’ internal state and ultimately influence movement decisions. Solid arrows show the flow of a physical process and dotted arrows are show the flow of information. b) The movement process is formulated on continuous time, where updating of the individual’s direction and behavioural state are independent processes. Behavioural state (Searching and Feeding) ultimately determines the processes used in selecting the next direction (represented by an angular probability density function of resource memory *g(θ)* and scent *d*_*i*_*(θ)* and the speed (*v*)/correlation (*τ*) values. The component processes of Memory (i.e., resource memory; blue) and Scent (red) in (a) are highlighted in the movement process model (b).

**Table 1.**
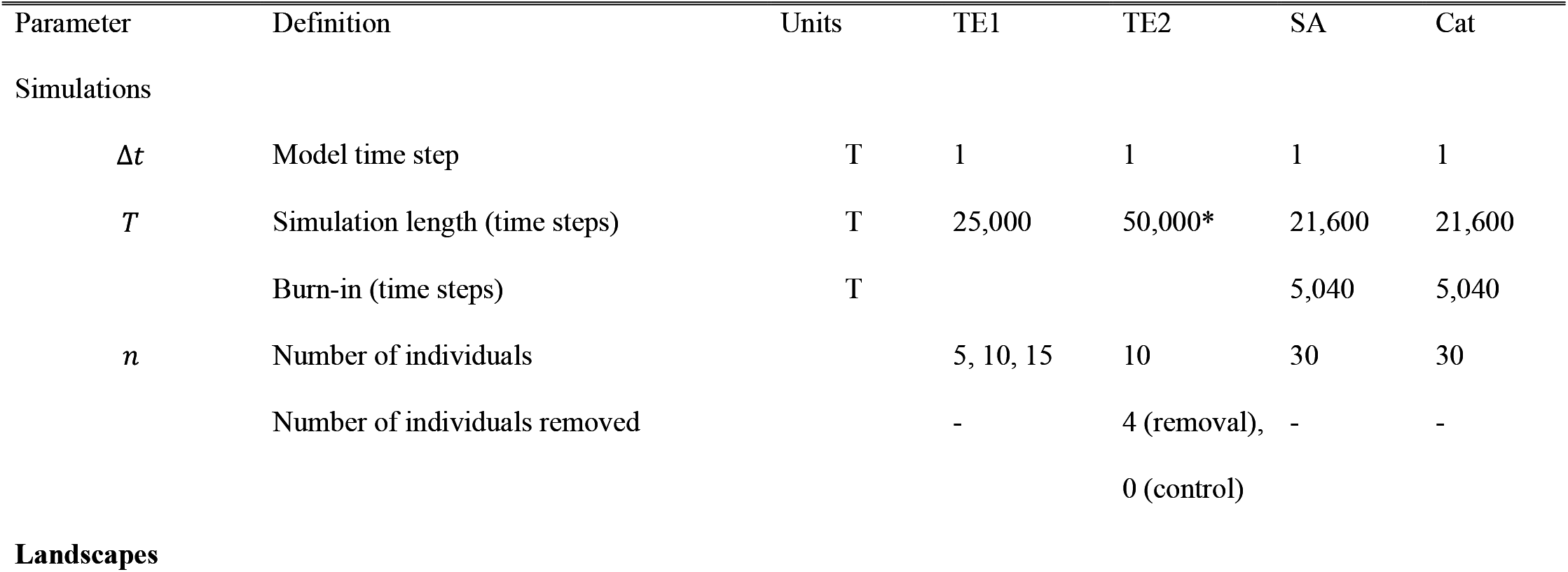
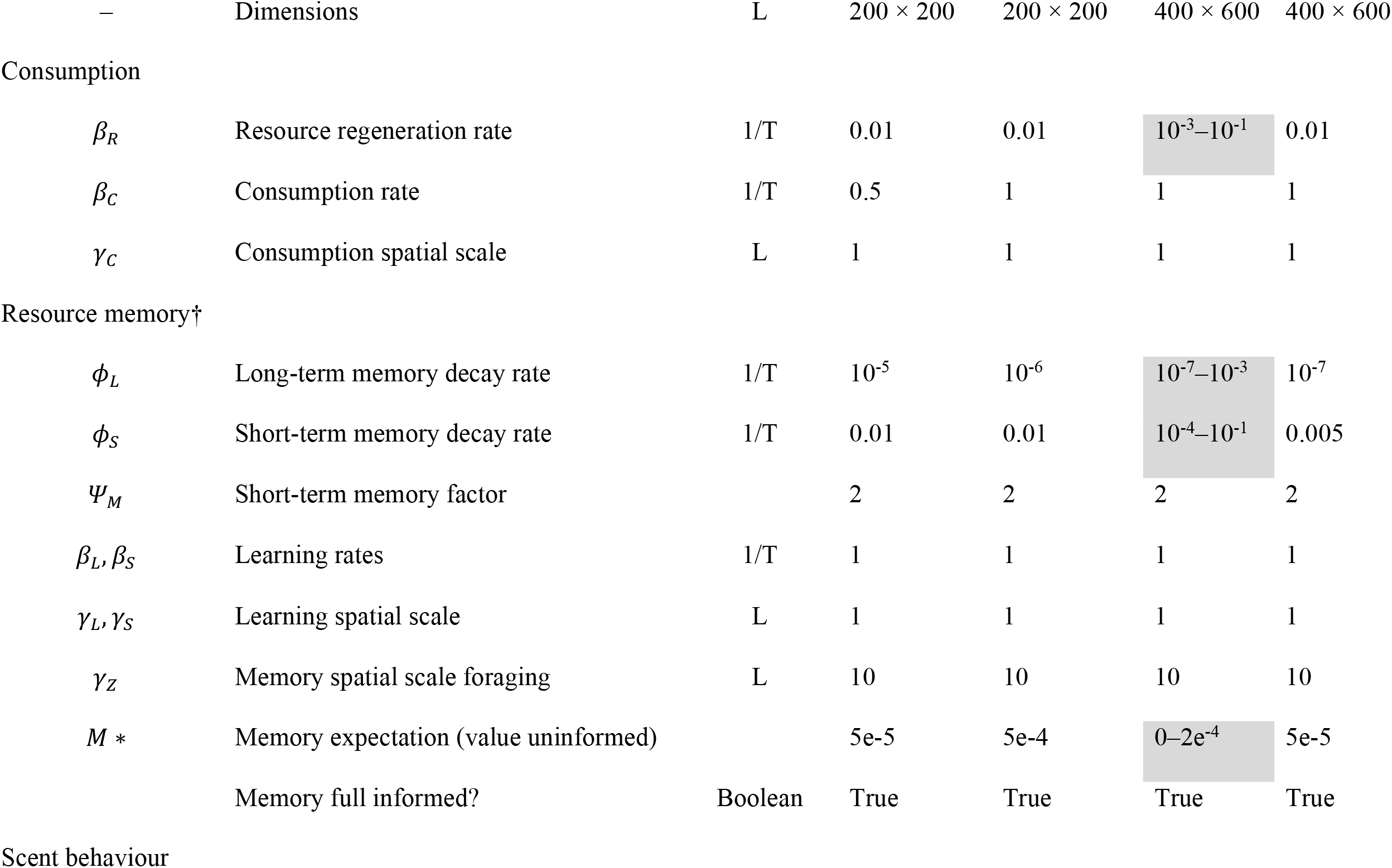
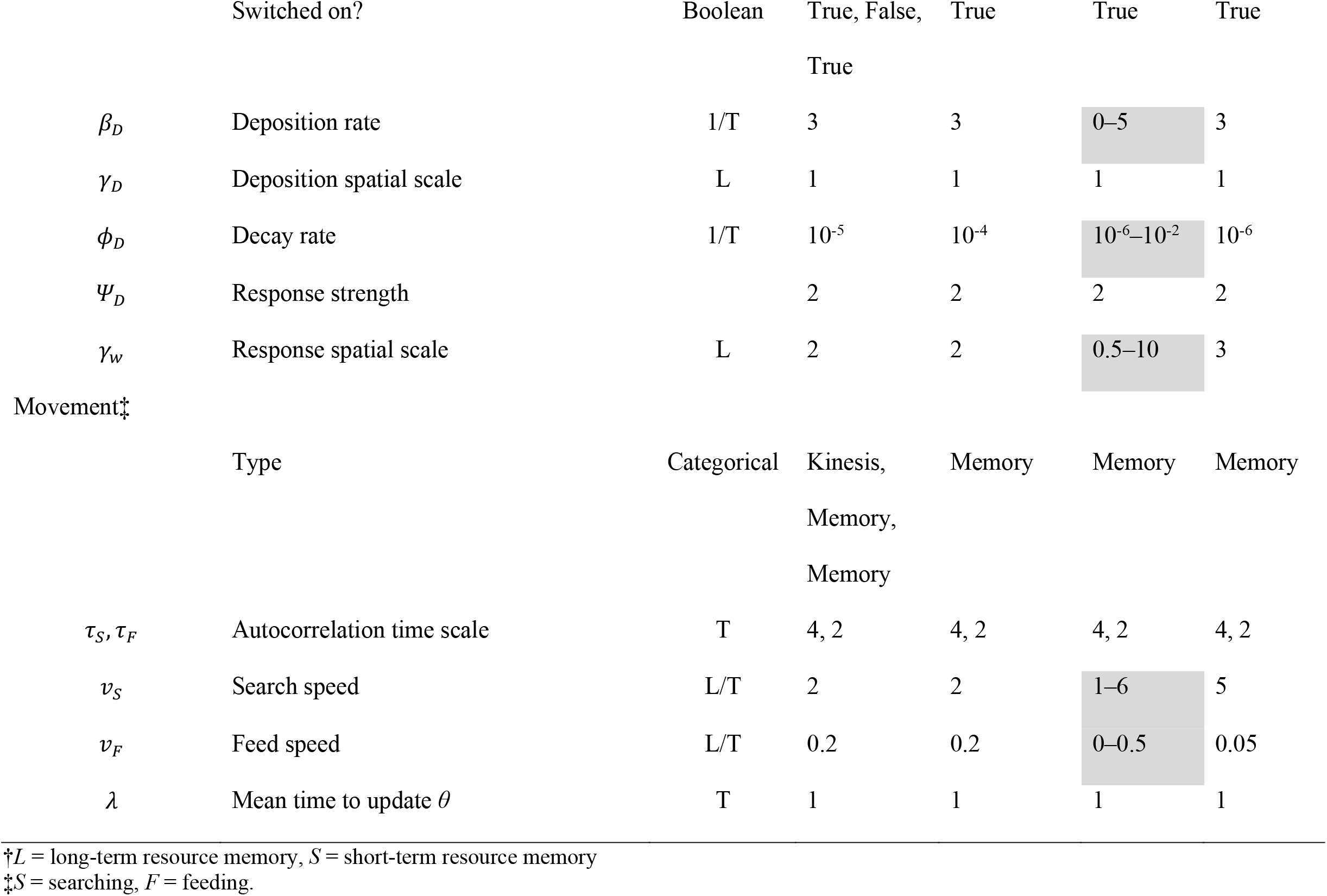
Overview of simulation parameters used in all four sets of simulations in the study. In theoretical example 1 (TE1), we tested the impacts of resource memory and territoriality on emergent home-ranges across three scenarios (territoriality only, resource memory only, territoriality and memory) and densities (*n* = 5, 10, 15). In theoretical example 2 (TE2), we test the impact of population manipulation by removing four out of ten individuals mid-simulation on emergent space use, and compare it with a control scenario without individual removal. In the sensitivity analysis (SA), we assessed the relative influenced of nine parameters across a range of values (grey cells). Finally, we ran simulations with a set of parameters calibrated to the empirical feral cat data (Cat). Units are arbitrary in the simulations, L is used for generic length units and T is used for generic time units.

### 2.1 Movement model description

Our model builds upon a two-state memory-based foraging model (Bracis *et al*. 2015) to include multiple individuals that interact through scent-based conspecific avoidance behaviour. Simulated individuals move around a dynamic resource landscape, learning about the intrinsic quality of the landscape as they consume resources (**Fig. 1a**).

#### 2.1.1 Movement process

An animal’s movements through the landscape are described by a continuous trajectory with a current position of 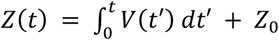, with a velocity of *V*(*t*) and initial position of *Z*_0_. The autocorrelated, directed, continuous movement process is given by

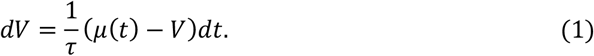

This is similar to the Ornstein-Uhlenbeck process where *τ* is the time scale of autocorrelation and instead of the white noise component, stochasticity is introduced through the bias vector *µ*(*t*) = (*v*,∠*θ*) of magnitude *v* = ||*µ*(*t*)|| (controlling average speed) and angle *θ* (direction). A Poisson process with rate parameter *λ* determines when angle *θ* is updated, which is then selected from an angular probability distribution derived from resource memory or scent processes, depending on the behavioural state. Finally, individuals switch between feeding and searching states, characterised by different values for *τ* and *v*, based on the current resource consumption *C*(*t*) (**Fig. 1b**).

#### 2.1.2 Resource memory

The resource *Q* is modelled as continuously varying in space across the landscape. Resources deplete as they are consumed by individuals and regenerate logistically at a rate of *β*_*R*_, but do not shift in space. Thus, it is advantageous for the individual to leave recently depleted patches but return to high quality areas over the long term. Animals consume resources according to a spatial kernel (a bivariate normal distribution with length scale *γ*_*C*_) and consumption rate *β*_*C*_. Animals have a resource memory with two different streams of information: a short-term stream *S* that pushes the animal away from recently visited locations even if they are attractive, and a long-term stream *L* that attracts the animal back to high quality areas (Van Moorter *et al*. 2009). The latter can either be initiated fully informed, with the intrinsic resource quality *Q*_0_, or naively, with a homogenous map of value *M \** indicating the animal’s expectation for unvisited areas which can be more optimistic or pessimistic and thus affect exploratory tendency (see Bracis & Wirsing, In review). *M \** is also the value to which long-term memory *L* decays. The two streams are combined into a single memory map *M*, which is used to inform the movement process.

The resource memory contribution to this angular probability distribution is computed by integrating transects of the resource memory map radiating out with radius *r* from the individual’s location with the memory value at each point weighted by distance,

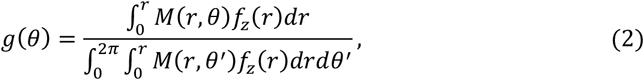

where *θ* represents the angle, and *f*_*z*_(*r*) is the kernel function that weights according to distance (here exponential with length scale *γ*_*z*_). The foraging memory movement model is described in further detail in Bracis *et al*. (2015).

#### 2.1.3 Scent-marking (territoriality)

As individuals move about the landscape, they also deposit scent, which decays over time, thereby marking their territory. The amount of scent on the landscape, *D*, is governed by the deposition rate, *β*_*D*_, (i.e., how much scent is deposited), the deposition spatial scale, *γ*_*D*_, (i.e, how broadly the scent is deposited in the vicinity of the forager) and the exponential decay rate *ϕ*_*D*_. *D* is limited to a maximum value of *D*0 and we take *D*0 to be one. Thus, the change in scent for each forager at location *z* = (*x, y*) is given by the equation

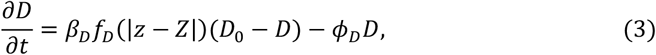

where *f*_*D*_ is the spatial kernel (here exponential with scale *γ*_*D*_). Scent deposition is tracked per individual, and foragers are indifferent to their own scent, but repulsed by the scents of all other conspecifics.

The repulsion of individuals by the scent of conspecifics is represented with an angular conspecific safety metric that scales between 0–1 but is not constrained to sum to 1 (like predation risk in Bracis *et al*. 2018b). This value represents the relative ‘safety’ in each direction in terms of avoiding conspecifics, with 0 meaning unsafe (high levels of conspecific scent) and 1 meaning safe (no conspecific scent). Safety is calculated by integrating the summed values of all other foragers’ deposited scent according to

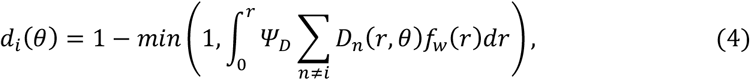

where *Ψ*_*D*_ is the response strength and*f*_*w*_(*r*) is a spatial kernel (here the exponential kernel with length scale *γ*_*w*_) that represents that decay of scent perception with increasing distance.

#### 2.1.4 Decision rules

To create the final angular probability distribution from which the angle *θ* is drawn to inform the movement process, the angular probability distribution based on the resource memory *g*(*θ*) is multiplied by the conspecific safety metric *d*_*i*_(*θ*) for individual *i*, then normalised, giving

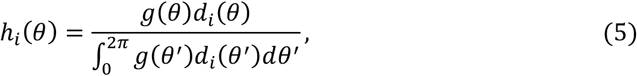

and the orientation angle of the bias term *θ*(*z*) = ∠*θ*(*z*), in the movement process is then drawn from *h*_*i*_(*θ*), which is specific for each individual.

### 2.2 Theoretical examples

The movement model was simulated on a 200×200 pixel heterogenous resource landscape, generated with the function *nlm_fbm* in R package *NLMR* (fractal dimension = 0.75). Initial locations of animals were random, and the landscape boundaries were reflective.

#### 2.2.1 Impacts of resource memory and territoriality

To demonstrate the effect of resource memory and territoriality on emergent HRs, we simulated animal movement trajectories for three scenarios: (1) territoriality only, (2) memory only; and (3) territoriality and memory in concert. In scenario 1, the movement process was a resource-dependent two-state random walk, with *g*(*θ*) in Eq. 5 being uniform while scenarios 2 and 3 use the memory process of Eq. 2. All three scenarios were run across three different densities of individuals (*n* = 5, 10, 15), creating 100 replicate simulations for each combination of scenario and density. Individual HR size was estimated from the simulation outputs using a kernel density estimator function *kernelUD* in the R package *adehabitatHR*. The 95% HR polygons were used to extract resource values within the landscape using the R packages *sp* and *raster*, after which the mean resource value within each HR was computed. Variance in each individual’s HR size was also visualised with respect to density and resource availability. A two-way ANOVA was used to evaluate the effect of density and scenario on log10-scaled HR size.

#### 2.2.2 Impacts of population manipulation

To demonstrate that HRs are both an emergent property and dynamic when conditions change in our model, we tested the effect of individual removal on remaining individuals’ HRs. We first simulated 100 datasets with a density of 10 individuals (i.e., a control setting). We simulated a further 100 datasets with the same parameters and starting conditions as the control setting, and then used a three-phase approach to remove individuals and quantify the effects of their removals on the HRs of the remaining individuals. The three phases were: (Phase 1) establishment of HR (*t* = 1–20,000); (Phase 2) remove four out of ten individuals (*t* = 20,001), followed by a transition period to allow the scent of removed individuals to decay to near zero (*t* = 20,001–30,000); and (Phase 3) exploration and establishment by remaining individuals (*t* = 30,001–50,000). To quantify the effects of removal, we compared the aggregate overlap in area (95% HR) between initial area covered by removed individuals (calculated from Phase 1 with 5,000 timesteps burn-in) and the area covered by remaining individuals before and after removal (calculated from Phase 1 and 3 respectively with 5,000 timesteps burn-in). A paired t-test was used to compare differences between the dependent before-after samples. For comparisons in the control setting, four of the 10 individuals were randomly “marked for removal” (but not actually removed). Aggregate overlap in area covered between “marked for removal” and “unmarked” individuals was then calculated using the function *over* from R package *rgeos*.

### 2.3 Model calibration to empirical data using sensitivity analysis

To facilitate a better mechanistic understanding of emergent properties of the model and to examine model application, we (1) conducted a sensitivity analysis (SA) to evaluate input parameter influence; and (2) used the SA results to guide calibration of parameters to match general patterns in an empirical case study that consisted GPS-locations from 11 adult female and male feral cats (*Felis catus*) from a 2019 eradication operation conducted on Kangaroo Island (South Australia) by the Department for Environment and Water South Australia (see **Text S1** for details on the empirical study and simulation setup).

#### 2.3.1 Sensitivity analysis

We assessed the relative influence of the following nine input parameters on emergent movement patterns: resource regeneration rate, searching speed, feeding speed, short-term memory decay rate, long-term memory decay rate, memory expectation, scent deposition rate, scent decay rate and scent response spatial scale. Emergent patterns were summarised with six summary statistics for 1,000 sampled input parameter combinations (one replicate simulation per combination): step-length, step-length distribution skewness, daily distance, HR size, HR size distribution skewness and HR overlap. The influence of input parameters on each respective summary statistic was estimated with boosted regression trees (BRT) to account for complex, non-linear relationships and parameter interactions (Elith *et al*. 2008; Prowse *et al*. 2016). We calculated the relative influence of each parameter (scaled to sum to 100), with higher numbers indicating stronger influence on the response. In models with interactions, we measured the interaction strength for top pairwise interactions between input parameters, with values near zero representing negligible interactions. Details on summary statistic calculation and model fitting are in **Text S2** and **S3** respectively.

#### 2.3.2 Parameter calibration to match empirical data

We proceeded with parameter calibration via a multi-step process: (1) optimising to a parameter set that best achieved the target values for the six summary statistics simultaneously (using respective BRT models); and (2) manual calibration to five target criteria: HR size and HR size skew (quantitatively evaluated), HR overlap, cumulative HR size and spatial distribution of locations (visually evaluated). The second step was iterated until no further improvement was found (i.e., improving one metric made another worse). We then ran 10 simulations using our final set of parameters and compared the outcomes with the empirical data. Details for this procedure are in **Text S4**.

All simulations (**Section 2.2** and **2.3**) were performed using Java v. 13.0.2 on a High-Performance Computing cluster. Analyses of model outputs were performed in R v. 3.6.1 (R Core Team 2020).

## 3 Results

### 3.1 Resource memory and territoriality lead to exclusive HRs shaped by resource quality and population density

Individuals in the territoriality-only scenario established distinct non-overlapping territories and space use was independent of resource (**Fig. 2a**). In the memory-only scenario, individual space use favoured high resource areas but overlapped substantially (**Fig. 2b**). With both territorial and resource memory processes operating in concert, individuals established distinct territories that utilised high resource areas with relatively little overlap (**Fig. 2c**).

**Figure 2.**
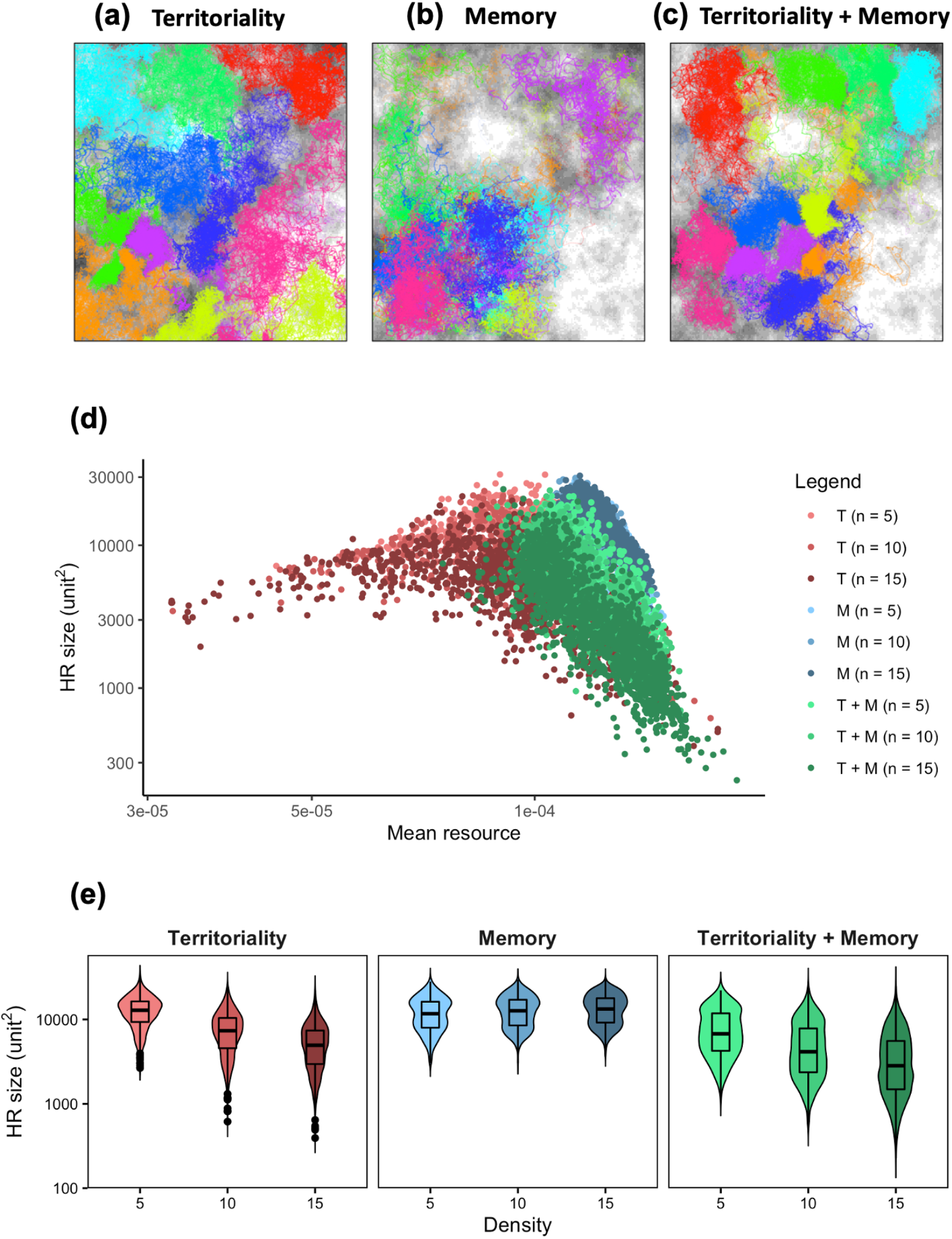
Simulation results of movement and home range from three scenarios (territoriality, memory, territoriality and memory): (a–c) sample realisations of animal movement trajectories (*n* = 10) from each scenario, where individual trajectories start at random (colours represent individuals); relationship between home range (HR) size and (d) mean resource value within HR (T: territoriality, M: memory, T + M: territoriality + memory) and (e) animal density (*n* = 5, 10, 15) from 100 replicates.

Across all three scenarios, individuals in low resource areas tended to have larger HRs than in high resource areas (**Fig. 2d**). This relationship was strongly log-linear in the memory-only and territoriality + memory scenarios (Pearson’s *r* = -0.75; *p* < 0.001), though variation in both HR size and mean resource were larger for the latter. The resource and HR size relationship in the territoriality-only scenario was consistent with the other scenarios at mid to high resource areas but starts to trend towards a positive HR-resource relationship in low resource areas, with a much smaller lower-limit of mean resource compared to the other two scenarios.

An increase in density caused a decrease in mean HR size in the territoriality-only (mean(HR size)_density=5/10/15_ = 11,860/6,670/4,600 unit^2^) and territoriality + memory (mean(HR size)_density=5/10/15_ = 6,730/4,200/2,840 unit^2^) scenarios, while it caused a slight increase in mean HR size in the memory-only scenario (mean(HR size)_density=5/10/15_ = 11,230/12,080/12,730 unit^2^; two-way ANOVA, *F*_2,8995_ = 450, *p* < 0.001; **Fig. 2e**). Overall, the smallest HRs were observed in the territoriality + memory scenario (two-way ANOVA, *F*_2,8995_ = 2,580, *p* < 0.001).

### 3.2 Individuals move into recently unoccupied areas

Aggregate HR overlap between remaining individuals and removed individuals was markedly higher in Phase 3 compared to Phase 1 (mean(HR size)_Phase 1/3_ = 7,000/14,430 unit^2^; paired t-test: *t*_99_ = -19.9, *p* < 0.001) of the removal simulations, while there was a smaller increase in the overlap between individuals (mean(HR size)_Phase 1/3_ = 6,580/8,980 unit^2^; paired t-test, *t*_99_ = -8.2, *p* < 0.001) in the control simulations (**Fig. 3**). One example removal simulation showed space use shifts in remaining individuals into areas previously occupied by removed individuals. Individuals in the control example demonstrated smaller shifts in space use on average, explaining the small increase in aggregate HR overlap from Phase 1 to 3.

**Figure 3.**
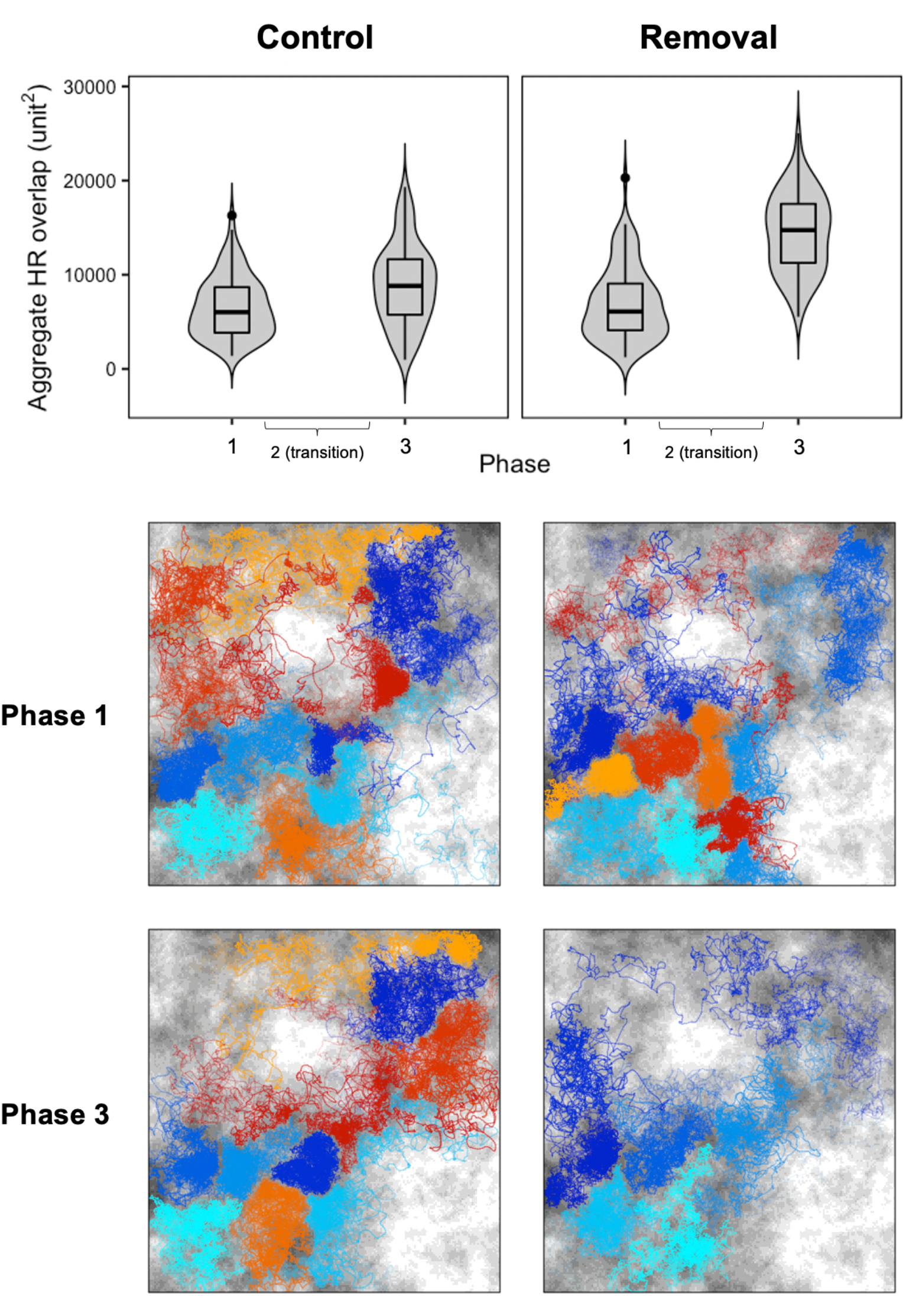
Aggregate home-range (HR) overlap between removed and remaining individuals before (Phase 1; *t* = 5,001–20,000) and after removal (Phase 3; *t* = 35,001–50,000) in the control and removal scenarios (100 replicates each). Beneath each scenario is a sample realisation of movement trajectories in Phase 1 (top panel) and 3 (bottom panel) of the control (left panel) and removal (right panel) simulation. Individual trajectories decrease in opacity according to time. Cool colours represent remaining individuals, warm colours represent removed individuals (in the control scenario, individuals were randomly grouped as remaining or removed for a comparison to be made).

### 3.3 Sensitivity analysis

HR size was strongly influenced by memory decay rates and scent decay rate (**Fig. 4a**). Short-term memory decay rate had a strong negative effect on HR size, whereas long-term memory and scent decay rates had strong threshold effects from around the mid-points of their parameter ranges, which increased constantly for long-term memory decay but tended to plateau at higher rates of scent decay (**Fig. 4b**). Interactions led to a combined effect that if scent decay rate was high, and either short- or long-term memory decay rate was low or high respectively, a 4–6 times increase in HR size was observed (**Fig. S7**). Longer right-tailed skews in HR size distribution were primarily caused by slower rates of scent decay, faster rates of short-term memory decay and larger scent response spatial scales (**Fig. 4c**). Smaller overlaps in HR were predominantly associated with slow scent decay rates and fast short-term memory decay rates (**Fig. 4d**).

**Figure 4.**
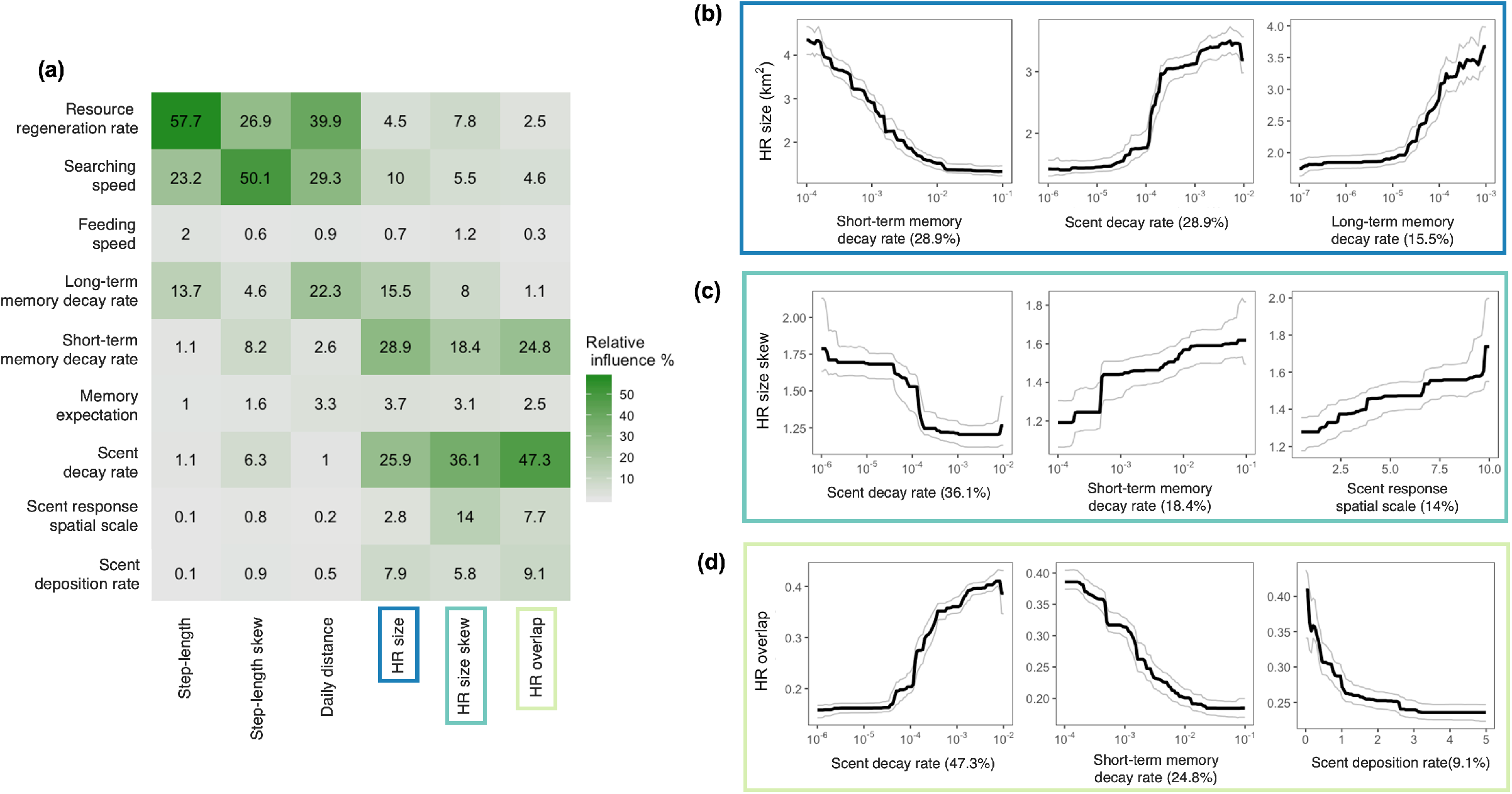
Results from the sensitivity analysis. (a) Relative influence (%) of nine model input parameters (rows) on six emergent movement summary statistics (columns) from a boosted regression tree (BRT) analysis. Partial dependency plots showing effects of the top three contributing input parameters on (b) home-range (HR) size, (c) home-range size skew and (d) home-range overlap. Partial dependency plots show the effect of a given explanatory variable on each home-range metric, while holding the effects of other explanatory variables at their average (black lines). We used 500 bootstrap replicates to calculate the 95% confidence intervals, depicted in grey. Note that some of the x-axes are on a log_10_ scale. Full plots showing partial effects of all parameters on each summary statistic can be found in **Fig. S3– S9**.

Distance metrics (step-length, step-length skewness, daily distance) were mainly influenced by resource regeneration rate and search speed (**Fig. 4a**). Step-length and daily distance increased, while step-length skewness decreased, with lower resource regeneration rates and higher search speeds (**Fig. S3**–**5**).

### 3.4 Empirical application

The simulated data generally captured the long right-tailed distribution of HR sizes that reflected populations of predominantly resident individuals (with smaller and distinct HRs), and a small number of transient individuals (with larger and less distinct HRs, and sparser location density) observed from the empirical cat movement data (**Fig. 5a**). Median HR was the same in the simulated and empirical data (1.9 km^2^). However, simulated HR size range tended to be smaller than the empirical data (0.5 vs. 1.3 km^2^ respectively) – approximately a tenth of simulated individuals had HRs smaller than one km^2^ and few ranged wider than nine km^2^ across the replicates. Furthermore, the cumulative HR in the simulations reflected a less stable system compared to the empirical data (i.e., curves did not approach a plateau for most individuals; **Fig. 5b**; **Table S2** for all). Spatial patterns of simulated locations resembled empirical movement patterns in that locations tended to cluster linearly along high resources, but to a markedly lesser extent. The intense clustering of locations in specific sites observed for cats was not reproduced in the simulated data (**Fig. 5c**).

**Figure 5.**
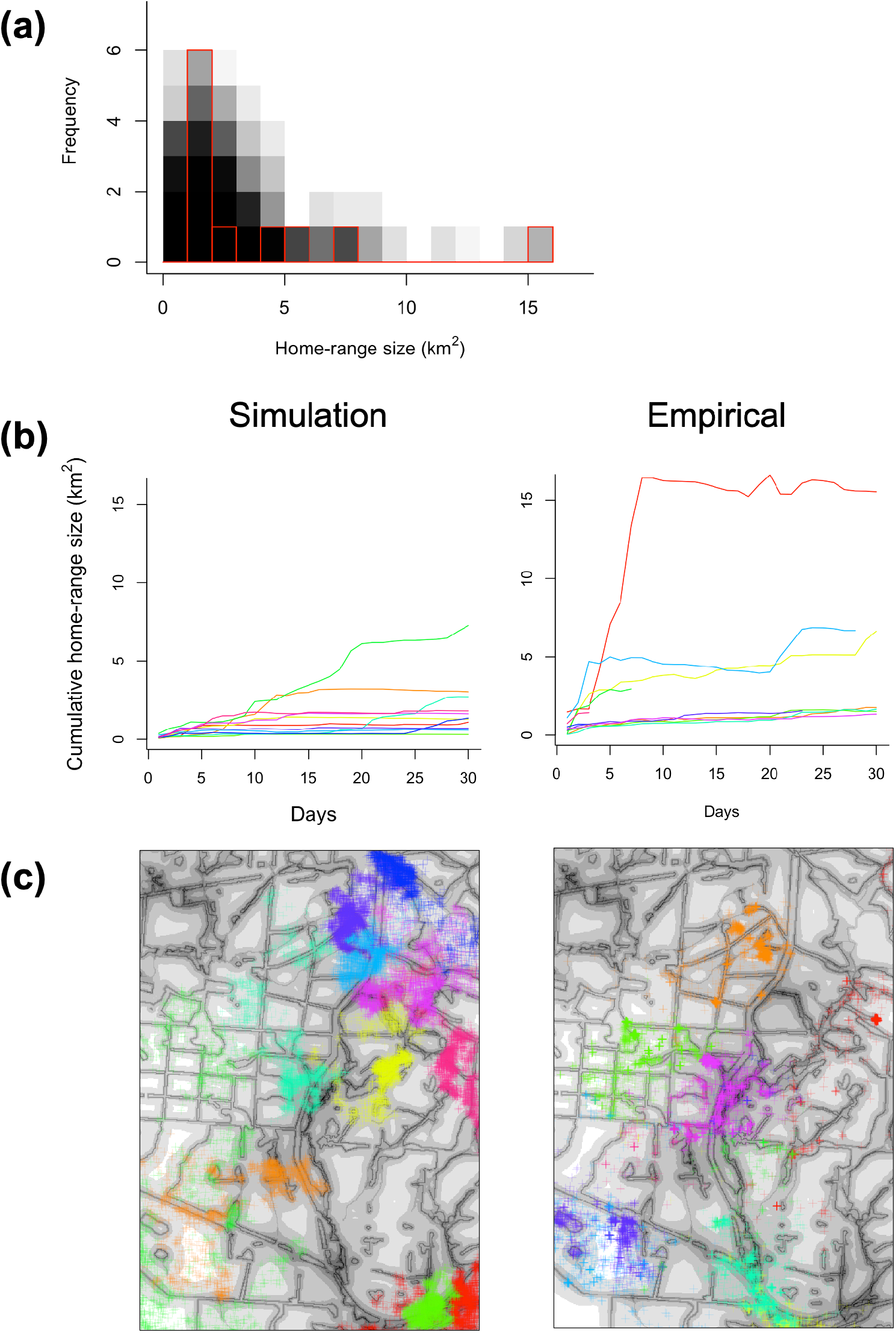
A comparison of movement patterns from parameterised simulations and the empirical data: (a) frequency distribution of home-range (HR) size of simulated individuals obtained from 10 replicates bootstrap sampled 10 times each (samples represented as 100 histograms in translucent black fill) to match the empirical sample of 11 tagged feral cats in Kangaroo Island, South Australia (histogram with empty fill and red border); (b) one realisation of cumulative HR size over time of 11 randomly selected individuals (left; individuals shown in different colours) against that of the observed feral cat ranges (right); and (c) their corresponding simulated movement locations (left) vs. observed feral cat movement locations (right; note that some cat locations go beyond the landscape shown here).

## 4 Discussion

HR formation is a ubiquitous feature of animal behaviour, yet we still lack a unifying theory explaining its emergence and the wide array of complex, dynamic spatio-temporal space use patterns observed in populations. Theoretical models have separately shown that memory-based resource use (Van Moorter *et al*. 2009; Bracis *et al*. 2015), nonterritorial conspecific avoidance (Riotte-Lambert *et al*. 2015), scent-based conspecific avoidance (Potts *et al*. 2012), and memory-based conspecific avoidance based on past interactions (Potts & Lewis 2016) can lead to the emergence of HRs. We integrated multiple drivers of space use, memory-based resource use and scent-based conspecific avoidance, to illustrate the emergence of exclusive HRs whose sizes were negatively related to resource and population density, and responsive to a changing conspecific environment, consistent with empirical observations. Neither mechanism completely controlled HR dynamics and their interactions strongly determined outcomes. The model can be applied to reproduce general spatial measures and patterns of feral cat space use, including the phenomenon of transience.

In our multi-individual simulation environment, two mechanisms were required to replicate patterns of HR formation in territorial animals: one that tended towards fidelity to previously visited, high quality areas (resource memory); and another that drove spatial avoidance between individuals (territoriality). The model captured realistic responses to density with the emergence of transient behaviour by some individuals as density increased and all high-quality areas were occupied. The absence of territoriality meant that individuals acted independently of conspecifics, which caused large overlaps in emergent HRs (**Fig. 2b**). The absence of resource memory resulted in unrealistic HR formation that was independent of resource quality; movement was only constrained by the scent of surrounding conspecifics (**Fig. 2a**). The combination of both mechanisms led to the emergence of distinct individual HRs distributed across high resource areas of the landscape (**Fig. 2c**). In addition to being spatially realistic, biologically meaningful relationships were also captured. Simulated individuals’ HR sizes correlated negatively with both resource availability and population density, which is congruent with ecological evidence from both field and experimental studies (Baker *et al*. 2000; Santangeli *et al*. 2012; Šálek *et al*. 2014; Schoepf *et al*. 2015). This is biologically intuitive. Namely, when resources are abundant, an individual needs less space to meet its metabolic needs (McLoughlin & Ferguson 2016); when population density is high, the amount of available, unoccupied space is smaller, which limits individual HR sizes (Schradin *et al*. 2010).

Home-ranging is a dynamic property emerging from our model, continuously adapting to the dynamic resource landscape and presence of surrounding conspecifics. This was emphasised in our removal simulations, where remaining individuals migrated into areas previously occupied by removed individuals once their scent decayed. The phenomenon of expanding into newly vacated habitat can be found in numerous species, such as mice (Schoepf *et al*. 2015), chipmunks, (Mares *et al*. 1976), coyotes (Moorcroft *et al*. 2006), and red foxes (Potts *et al*. 2013). Non-mechanistic models (e.g., with imposed HR centres and/or boundaries) fail to respond to changing environments (e.g., individuals leaving or dying, fluctuating resource availability), whereas in our model dynamic behaviour arises directly from underlying mechanisms. Notably, the memory mechanism in our model provides the basis for HR stabilisation in the absence of territoriality, which existing mechanistic territorial models either lacked or imposed through the assignment of HR centres (Potts & Lewis 2014). Moreover, the ability to control the relative strengths of each mechanistic component allows modulation of remaining individuals’ spatio-temporal responses to vacancies, which can vary according to sociodemographic factors (Frank *et al*. 2018).

The sensitivity analysis revealed that emergent HR sizes were most sensitive to memory (short- and long-term) and scent decay rates. These rate parameters essentially control the decay in effects (i.e., duration of time-scale of effects) of modelled mechanisms that act on space use. Short-term memory is a repulsive force pushing the individual to move beyond areas it has visited, where a slower decay results in more exploratory (or diffusive) movement leading to larger HR sizes. On the other hand, long-term memory is an attractive force that attracts the individual to areas, based on either an *informed* memory of previously visited high resource areas, or an *uninformed* expectation that areas beyond the local area of movement (that are unexplored or forgotten) are better. If the animal’s expectation for unexplored area matches the mean resource value, faster long-term memory decay rates led to larger HR sizes because the memory expectation was generally better than most individuals’ local area of movement (i.e., resource distribution was skewed so the mean was greater than the median value). If unexplored area expectations were low, we expect long-term memory decay to have the opposite negative effect on HR size. Unlike resource memory, the effect of scent is externally conferred by conspecifics. When conspecific scent lingers at slower decay rates, it acts as an external negative pressure limiting HR sizes. While most individuals have smaller HR sizes with slow scent decay rates, a few individuals ranged widely in low resource areas, causing the longer right-tailed skew in overall HR size distribution.

The effects of resource memory and scent on emergent HRs from our model raises an interesting point: HR formation in territorial animals can be attributed to both memory-mediated philopatric behaviour operating at the individual-level (i.e., inherent exploratory vs. recursive tendency) and spatial competition at the population-level. Existing theoretical models have independently demonstrated that distinct and stable HRs can form from either resource memory-mediated movement (Van Moorter *et al*. 2009; Bracis *et al*. 2015) *or* scent- or memory-mediated conspecific avoidance (Potts *et al*. 2012; Giuggioli *et al*. 2013; Ellison *et al*. 2020) alone. In territorial species, both mechanisms are likely crucial to HR formation. While site revisits owing to memory processes are commonly observed (Bracis *et al*. 2018a), it is also common for animals to ‘probe’ the boundaries of their HR and expand into newly unoccupied habitat. Thus, to consider the degree to which each component drives HR formation, it is important to combine them within the same modelling framework to understand how territorial animals integrate the opposing motivators and the relative strengths of each response, which is possible with different component parameterisations in our model. Mechanistic home-range analyses (MHRAs) on carnivores have integrated and disentangled the effects of scent-mediated conspecific avoidance from other drivers of HR formation (e.g., resource selection and steep terrain avoidance) in their modelling frameworks (Moorcroft *et al*. 2006; Bateman *et al*. 2015). However, the use of memory as mechanistic basis for philopatry (e.g., site fidelity) within analytical models is still in its’ infancy. The analytical framework used in recent studies to incorporate memory-mediated behaviours (e.g., step selection and conspecific avoidance, Oliveira-Santos *et al*. 2016; Ellison *et al*. 2020) can plausibly be extended to memory-mediated philopatry. Integrating both territoriality and memory-mediated philopatry into MHRAs to uncover key drivers of HR formation is an important area of future work (Fagan *et al*. 2013).

Despite challenges in simultaneously fitting multiple metrics to a real system with our movement model, we successfully approximated the general patterns. The BRT models of the sensitivity analysis output were able to approximate simulation outcomes and facilitated parameter calibration to real data using multiple summary metrics. Following manual calibration, our simulations reproduced the target median and positively skewed variation in HR size of the empirical cat GPS dataset. The underestimate of the upper limit of HR size may be due to the spatial limitation of the truncated simulation landscape (to prevent long simulation times) from the total landscape covered by the tagged feral cats. Nonetheless, we were able to replicate the emergence of wide-ranging individuals (or transients) amidst resident individuals, as seen by the positively skewed variation in individual HR size, reflecting a real phenomenon observed in our data and many other territorial species (Holmes *et al*. 1996; Mitchell *et al*. 2015; Hinton *et al*. 2016). While transience itself is a complex movement phenomenon, its emergence can occur with the core mechanisms of resource memory and territoriality, without modelling further complexities such as individual differences in motivation (e.g., varying exploratory tendencies).

We were only able to partially replicate finer scale spatio-temporal patterns estimated for feral cats (i.e., cumulative HR) for several possible reasons. First, the less stable simulated cumulative HR curves could indicate that our burn-in period was not long enough for individuals to establish stable HRs and that burn-in time should be considered in parameter calibration. However, the simulated patterns were not unrealistic and have been observed in real systems (e.g., Eurasian lynx, Breitenmoser-Würsten *et al*. 2007); Brazilian mesocarnivores, Bianchi *et al*. 2016). Secondly, we were not able to quantitatively describe the cumulative HR curves. Quantitative characterisation of such curves (including similar metrics such as net squared displacement) is a challenging and active area of research (Spitz *et al*. 2017).

Ultimately, improving resemblance to our real data likely requires additional mechanisms for movement behaviours common in carnivores. For example, the lack of location point clusters and concentrations along linear resources in our simulated data suggests that we did not account for intermittent resting behaviour and preferential use of corridors respectively (Noss *et al*. 1996). Integrating rest site and corridor layers into the resource map could potentially improve emergent spatial patterns. However, the underlying utilisation mechanism is different from modelled behaviours (of continuous resource consumption), which may warrant building additional mechanisms and landscape layers into the model (e.g., a resting state dependent on the animal’s internal bioenergetic state; landscape permeability). Additionally, methods that attempt to identify site fidelity and quantify recursions can potentially be used to parameterise such a model (Richardson *et al*. 2017; Bracis *et al*. 2018a).

Our flexible modelling framework simulates continuous-time movement trajectories that can be applied to a variety of populations exhibiting cue-based territorial behaviour. Models can be calibrated and validated against empirical movement data, which is becoming increasingly available (Kays *et al*. 2015). While further development may be necessary to replicate some fine-scale movement patterns, our model can be used to simulate generic memory-informed and territorial space use, in terms of broadly matching HR characteristics. Potential applications include simulation-based evaluations of behavioural structure in animal movement (e.g., Gurarie *et al*. 2016) and population estimation of mobile animals (e.g., Theng *et al*., In Review). Another potential application is in animal-mediated seed dispersal research, which has identified the need to integrate frugivory and disperser movement and to consider more process-based movement approaches (Nield *et al*. 2019).

In conclusion, we developed a broadly applicable modelling framework that integrates two key underlying mechanisms of home-range formation, resource memory and territoriality, to simulate complex patterns of territorial animal space use. Our model application to a target population of feral cats demonstrated that general space use patterns evident in the empirical data could be approximated through simulation, and while replication of finer-scale space use patterns is plausible, we have identified key considerations for future investigations. Our understanding of how animals integrate multiple aspects of spatial localisation is only beginning to emerge, and our modelling approach can provide a foundation for sophisticated theoretical models of space use in interacting animals.

## Acknowledgements

M.T. was supported by an Adelaide Postgraduate Scholarship Award (International). This work was supported with supercomputing resources provided by the Phoenix HPC service at the University of Adelaide. Empirical data was from a 2019 feral cat control study conducted by the Department for Environment and Water South Australia. The authors acknowledge the Indigenous Traditional Owners of the land on which the University of Adelaide is built – the Kaurna people of the Adelaide Plains.

