## Supplementary Information for "The integrated role of resource memory and scent-based territoriality in the emergence of home-ranges" (from another version of this preprint: version 2, 2021-07-14)

**Text S1.** Description of the empirical data and parameterisation of the simulation landscape
**Text S2.** Sensitivity analysis details
**Text S3.** BRT model fitting and selection
**Text S4.** Parameter calibration details
**Table S1.** Comparison of output metrics from empirical data, predictions from BRT models and simulations (i.e., realised data).
**Table S2.** Results from each step of parameter calibration.
**Figure S1.** Feral cat locations.
**Figure S2.** Estimated prediction error curves and their standard errors for boosted regression tree (BRT) models of each summary statistic.
**Figure S3.** Partial effects of input parameters on step-length
**Figure S4.** Partial effects of input parameters on step-length skewness
**Figure S5.** Partial effects of input parameters on daily distance
**Figure S6.** Partial effects of input parameters on home-range (HR) size
**Figure S7.** Pairwise parameter interactions explaining home-range (HR) size
**Figure S8.** Partial effects of input parameters on home-range (HR) size skewness
**Figure S9.** Partial effects of input parameters on home-range (HR) overlap

**Text S1. Description of the empirical data and parameterisation of the simulation landscape**

We used data from 11 GPS-tagged adult female and male feral cats from a 2019 feral cat control study conducted in Kangaroo Island (South Australia) by the Department for Environment and Water South Australia (obtained with permission; **Fig. S2**). Location data for all individuals started on the same date (13 February 2019), and fixes were taken every 15 minutes between 6 pm – 6 am (active period) up until mortality from poison or collars falling off.

As we aimed to relate simulated data to the real dataset, we used a simulation resource landscape representative of the 4 × 6 km study area (spatial extent of [763421, 767421] to [6029884, 6035884]; UTM zone 53H). One spatial unit of our simulation landscape was equivalent to 10 m. Resource values were derived by calculating the 1–($\log_2$+1) scaled Euclidean distance from three environmental variables hypothesised to be of importance to cats – water courses, vegetation and buildings (J. Smith, personal communication, 2 March 2021) – expressing non-linear preference for areas nearer these resources compared to areas further from them. Environmental layers were obtained from https://data.environment.sa.gov.au/.

The number of simulated individuals was kept constant at 30, loosely based on the estimated density of cats in the study area (Berris *et al.* 2019). Since territorial behaviour (i.e., conspecific avoidance) was mostly exhibited within the same sex (**Fig. S2**), we attempted to parameterise our simulations to a single sex. This translated to a density of 1.25/km$^2$ in our simulation landscape, which is slightly less than half of the empirical density estimate of 2.98/km$^2$.

Each simulation was run for 21,600 timesteps with a prior burn-in of 5,040 timesteps. We treated the output as equivalent to 30 days (i.e., one timestep as one minute on a 12-hour activity day).

**Text S2. Sensitivity analysis details**

***Parameters of interest.*** Since we were interested in the physical properties of emergent movement patterns, we selected parameters that were hypothesised to be of significant influence: resource regeneration rate, searching speed, feeding speed, short-term memory decay rate, long-term memory decay rate, memory expectation, scent deposition rate, scent decay rate and scent response spatial scale. Length parameters (searching and feeding speed), resource regeneration and consumption rates were hypothesised to directly influence distance metrics (step-length, step-length distribution skewness, daily distance covered). We selected two memory parameters, the short- and long-term decay rate, that were predicted to affect the exploratory tendencies of individuals via repulsion (from recently visited areas) and attraction (to long-term memory of resource quality) respectively. As the role of scent was postulated to play an important role in mediating HR size and degree of HR overlap, we selected three of five scent parameters for the SA (scent deposition and decay rate, scent response strength).

***Summary statistics.*** Summary statistics for each replicate simulation were calculated from data subsampled at 15-minute intervals (one timestep = one minute) to match the temporal resolution of the cat data and were calculated as follows:

**Step-length** was measured as the Euclidean distance between consecutive steps, and the median of all individuals' median step-length was taken as the summary statistic for the replicate.

**Step-length distribution skewness** was the replicate median skewness measure across all individuals' step-length distributions.

**Daily distance** was measured as the sum of all step-lengths within each 12-hour bloc (treated as one day), and the median of all individuals' median daily distance covered was taken as the summary statistic for the replicate.

**HR size** was the replicate median of individuals' 95% kernel density estimate of home-range area.

**HR size skewness** was the skewness measure across individuals' HR size.

**HR overlap** was the median across individuals' maximum pairwise (proportional) overlap in HR size.

For the first three summary statistics, we first converted the data to an *ltraj* object (of R package *adehabitatLT*), which automatically calculates the Euclidean distance between each consecutive step for each individual. HR size was calculated using the kernel density estimator function *kernelUD* in the R package *adehabitatHR*. For HR overlap, we calculated each individual's pairwise proportional overlap in HR with every other individual, which was taken as the overlapped area (using function *gIntersection* and *gAre*a of R package *rgeos*) divided by the pair's merged HR area (using function *gUnion* and *gAre*a of R package *rgeos*). Since most individual pairs are not adjacent and hence do not overlap, we prevent the effect of zero-inflation by taking the maximum value for each individual (i.e., the conspecific that it overlaps with the most). We used a proportional measure of HR overlap rather than an absolute measure to account for the confounding effect of varying HR sizes.

### Text S3. BRT model fitting and selection

To optimise sampling within a high dimensional parameter space, we drew 1,000 Latin hypercube (LH) samples of parameter combinations uniformly distributed over ranges given by **Table 1** with the function *randomLHS* in the R package *lhs*. LH sampling implements an equal-area subdivision of the sample space and then samples randomly within each subdivision (McKay *et al.* 1979). We ran movement simulations for 1,000 LH samples, one sample (replicate) per parameter combination. We traded-off more replicate runs per sample to more extensively sample across parameter space as it more accurately captures the input-output relationship of the simulation model (Prowse *et al.* 2016).

The influence of input parameters on each respective output metric (i.e., response) was estimated with boosted regression trees (BRT) to account for complex, non-linear relationships and parameter interactions (Elith *et al.* 2008; Prowse *et al.* 2016). We fitted multiple models for each response by varying the BRT interaction depth (or *tree complexity*) permitted. We fit BRT models with the R package *dismo* using the function *gbm.step* with a learning rate of 0.01, a bag fraction of 0.75, and a tree complexity of 1–9, and optimised the number of fitted trees based on fivefold cross-validation (CV). We assumed Gaussian errors for all the response variables. We selected the least complex model (lowest interaction depth) within one standard error of the best model (lowest CV deviance) (see Section 7.10 in Hastie *et al.* (2013)). If CV deviance for a response was less than 0.01 at tree complexity (*tc*) of 1, we automatically selected this model.

We calculated the relative influence of each input parameter, calculated as the number of times the parameter is selected for splitting a tree, weighted by the improvement resulting from that split, averaged over all trees. It was then scaled to sum to 100, with higher numbers indicating stronger influence on the output metric. We then constructed partial dependency plots to visualise the relationship between each input parameter and output metric. In models with a *tc* > 1, we measured the interaction strength for top pairwise interactions between input parameters, with values near zero represent negligible interactions.

### Text S4. Parameter calibration details

***Coarse-tuning.*** We started the calibration process by first determining an optimised parameter set with which to achieve the target values for the six summary statistics simultaneously. Their respective BRT models were used to predict and optimise output to target statistics, using the base R function '*optim*'. Optimisation was initialised with appropriate starting values determined by the SA rather than random values because of convergence issues (e.g., local maxima, step-like predictions did not fit method's smooth functions). We eventually prioritised optimisation to HR statistics (size, size skewness, overlap) because the step-length and daily distance statistics' respective upper ranges (output from the SA) had little and no overlap with the target (i.e., empirical) values (did not emerge from the range of input values). We then ran 10 simulations with this parameter set.

***Manual fine-tuning.*** Since the simulation output's concordance with HR statistics and spatio-temporal patterns was low (**Table S2**), subjective decisions had to be made as we proceeded with a manual calibration. Here, we added two criteria (cumulative HR size and spatial distribution of locations) to supplement the reductive measure of HR size, which alone could not capture the complex spatio-temporal properties of animal trajectories. We visually compared these patterns (and did not include them in the SA) because we were unable to derive a quantitative metric (e.g., individual cumulative HR curves were unique, and no one function fit closely to individual curves). HR overlap was also evaluated visually because the empirical measure was based only on a subset of the individuals occurring in the area. Thus, our target criteria for the manual-tuning procedure consisted: HR size, HR size skew, HR overlap, cumulative HR size and spatial distribution of locations.

We evaluated the output's concordance with the five target criteria and manually adjusted the parameter set based on that information. This process was iterated until no further improvement was found (i.e., improving one metric made another worse). We then ran 10 simulations using our final set of parameters and compared the outcomes with the empirical data.

**Table S1.** Comparison of output metrics from empirical data, predictions from BRT models and simulations (i.e., realised data). The mean (within one standard deviations) of all 10 simulations is presented.

| **Metric** | **Empirical** | **Prediction** | **Realised (simulations)** |
|---|---|---|---|
| Step-length (km) | 0.07 | 0.05 | 0.06 ± 0.003 |
| Step-length skewness | 2.2 | 1.8 | 1.6 ± 0.06 |
| Daily distance covered (km) | 4.8 | 3.0 | 3.4 ± 0.17 |
| HR size ($km^2$) | 1.9 | 1.8 | 1.9 ± 0.28 |
| HR size skewness | 1.8 | 1.6 | 1.4 ± 0.72 |

**Table S2. Results from each step of parameter calibration.** The process started with a set of parameters optimised from initial values deemed appropriate by the sensitivity analysis. Subsequently, parameters were manually fine-tuned with the aim of improving concordance with the target metrics. This process was iterated until we achieved four out of five of the criteria. The output for HR size (median) and skewness are the mean result from 10 simulations, unless indicated with a * (where result was from one simulation).

| Iteration | Input | Output |
|---|---|---|
| **---- *Course-tuning* ---** |  |  |
| Optimised set (SAoptim01) | *Parameters* — *Values*<br>$\beta_R$ — $10^{-3}$<br>$\phi_L$ — $10^{-7}$<br>$\phi_S$ — $10^{-3}$<br>$M$* — $2e^{-4}$<br>$\phi_D$ — $10^{-6}$<br>$\beta_D$ — 3<br>$\gamma_W$ — 3<br>$v_S$ — 5<br>$v_F$ — 0.5 | *HR size (median)*: 1.4<br>*HR size skew*: 1.6<br>*HR overlap*: Yes<br><br>*Cumulative HR*:<br>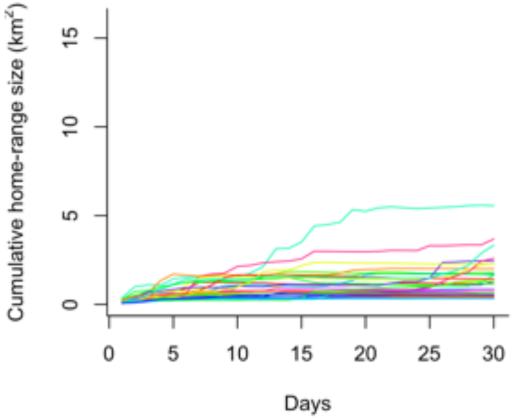<br>*Locations*:<br>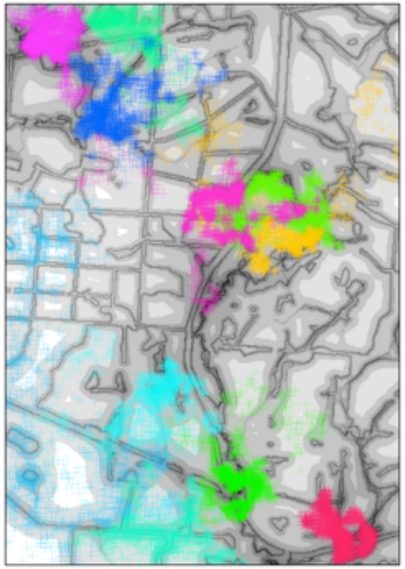 |
| **---- *Manual fine-tuning* ---** |  |  |
| 1 (SAoptim02) | *Parameters* — *Values*<br>$\beta_R$ — $10^{-3}$<br>$\phi_L$ — $10^{-7}$<br>$\phi_S$ — $10^{-3}$ | *HR size (median)*: 1.4<br>*HR size skew*: 1.6<br>*HR overlap*: Yes<br>*Locations*: |

| | | | |
|---|---|---|---|
| | $M^*$ 2e-4<br>$\phi_D$ $10^{-7}$<br>$\beta_D$ 2<br>$\gamma_w$ 3<br>$v_S$ 5<br>$v_F$ 0.5 | *Cumulative HR*:<br>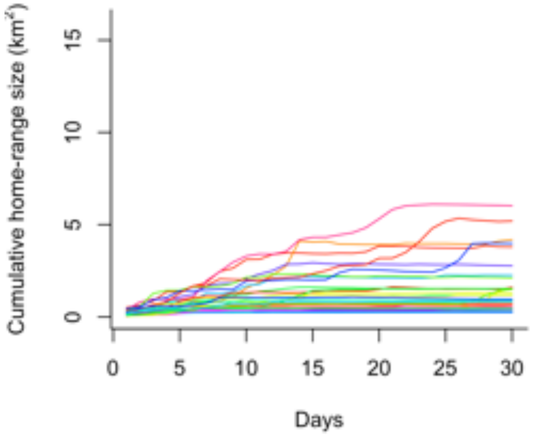 | 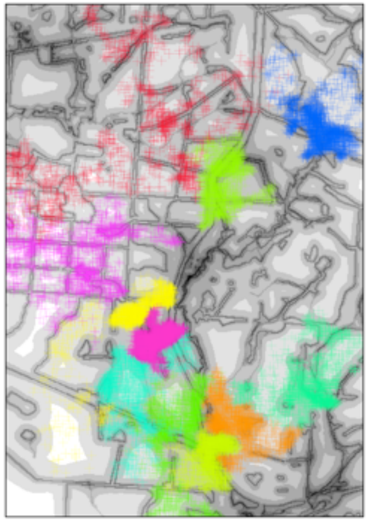 |
| 2<br>(SAoptim03) | *Parameters* *Values*<br>$\beta_R$ $10^{-3}$<br>$\phi_L$ $10^{-7}$<br>$\phi_S$ $10^{-3}$<br>$M^*$ $10^{-4}$<br>$\phi_D$ $10^{-6}$<br>$\beta_D$ 2<br>$\gamma_w$ 3<br>$v_S$ 6<br>$v_F$ 0.5 | *HR size (median)*: 1.9<br>*HR size skew*: 2.1<br>*HR overlap*: Yes<br><br>*Cumulative HR*:<br>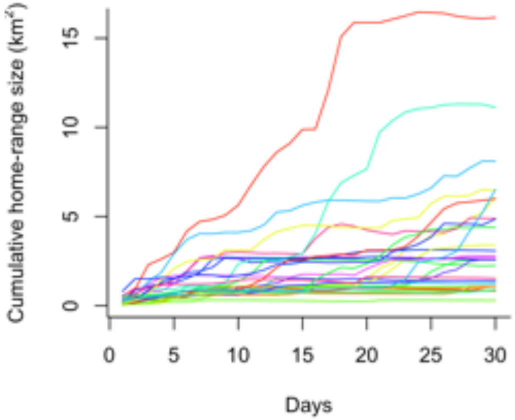 | *Locations*:<br>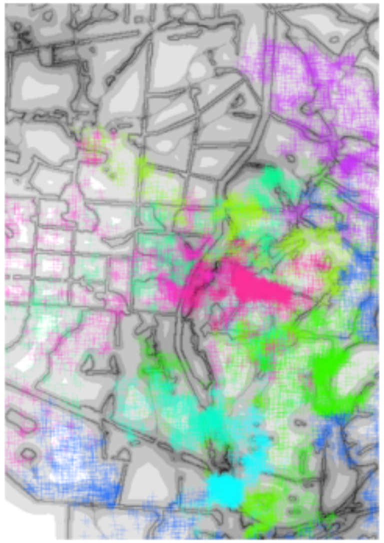 |

| | | | |
|---|---|---|---|
| 3<br>(SAoptim05) | *Parameters* *Values*<br>$\beta_R$ 5e$^{-3}$<br>$\phi_L$ 10$^{-4}$<br>$\phi_S$ 5e$^{-3}$<br>$M^*$ 5e$^{-5}$<br>$\phi_D$ 5e$^{-5}$<br>$\beta_D$ 2<br>$\gamma_W$ 3<br>$v_S$ 5<br>$v_F$ 0.05 | *HR size (median)*: 1.3<br>*HR size skew*: 1.7<br>*HR overlap*: Yes<br><br>*Cumulative HR*:<br>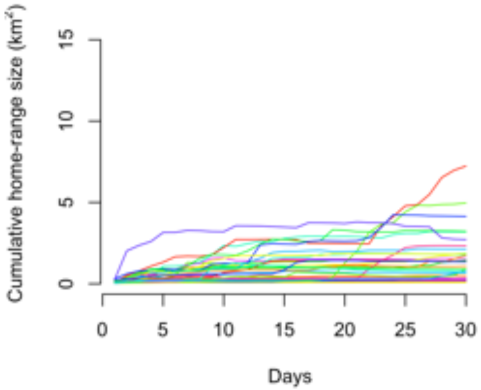 | *Locations*:<br>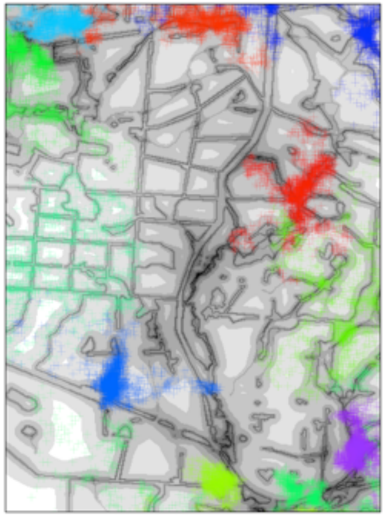 |
| 4<br>(SAoptim06) | *Parameters* *Values*<br>$\beta_R$ 2.5e$^{-3}$<br>$\phi_L$ 10$^{-4}$<br>$\phi_S$ 2.5e$^{-3}$<br>$M^*$ 5e$^{-5}$<br>$\phi_D$ 5e$^{-5}$<br>$\beta_D$ 2<br>$\gamma_W$ 3<br>$v_S$ 5<br>$v_F$ 0.05 | *HR size (median)*: 2.0<br>*HR size skew*: 1.6<br>*HR overlap*: Yes<br><br>*Cumulative HR*:<br>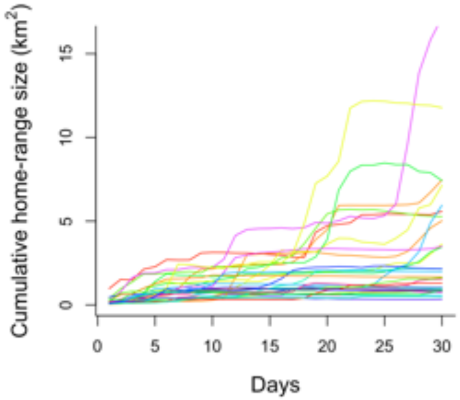 | *Locations*:<br>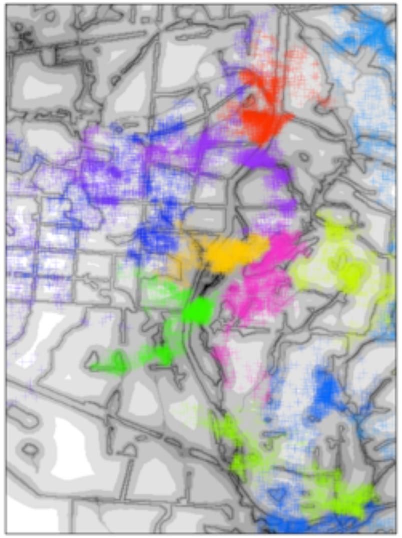 |

| | | | |
|---|---|---|---|
| 5<br>(SAoptim07)<br>* | *Parameters* *Values*<br>$\beta_R$ $10^{-3}$<br>$\phi_L$ $10^{-7}$<br>$\phi_S$ $10^{-3}$<br>$M^*$ $5e^{-5}$<br>$\phi_D$ $5e^{-5}$<br>$\beta_D$ 2<br>$\gamma_w$ 3<br>$v_S$ 5<br>$v_F$ 0.5 | *HR size (median)*: 2.0<br>*HR size skew*: 1.0<br>*HR overlap*: Yes<br><br>*Cumulative HR*:<br>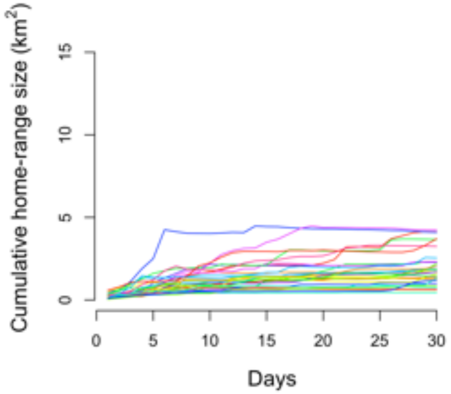 | *Locations*:<br>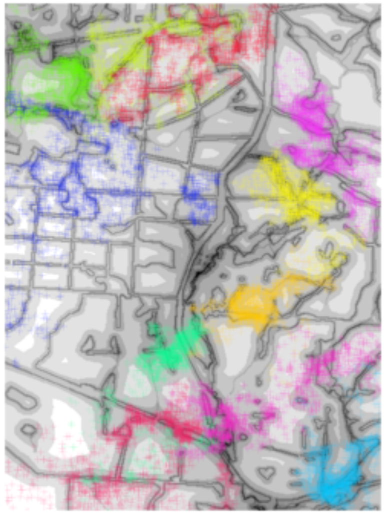 |
| 6<br>(SAoptim08) | *Parameters* *Values*<br>$\beta_R$ $10^{-3}$<br>$\phi_L$ $10^{-7}$<br>$\phi_S$ $10^{-3}$<br>$M^*$ $5e^{-5}$<br>$\phi_D$ $10^{-6}$<br>$\beta_D$ 2<br>$\gamma_w$ 3<br>$v_S$ 5<br>$v_F$ 0.5 | *HR size (median)*: 1.9<br>*HR size skew*: 1.4<br>*HR overlap*: Yes<br><br>*Cumulative HR*:<br>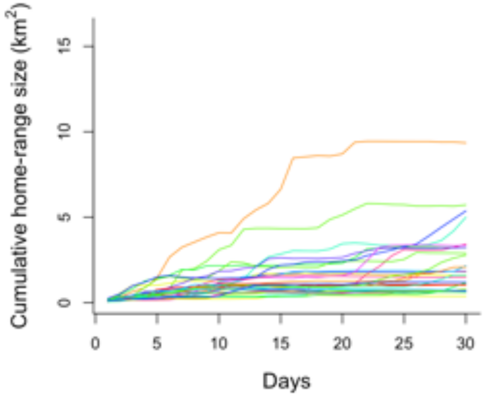 | *Locations*:<br>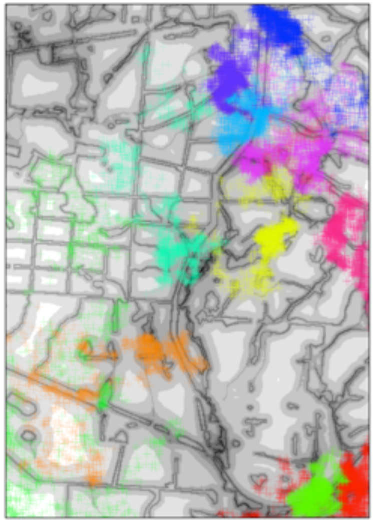 |

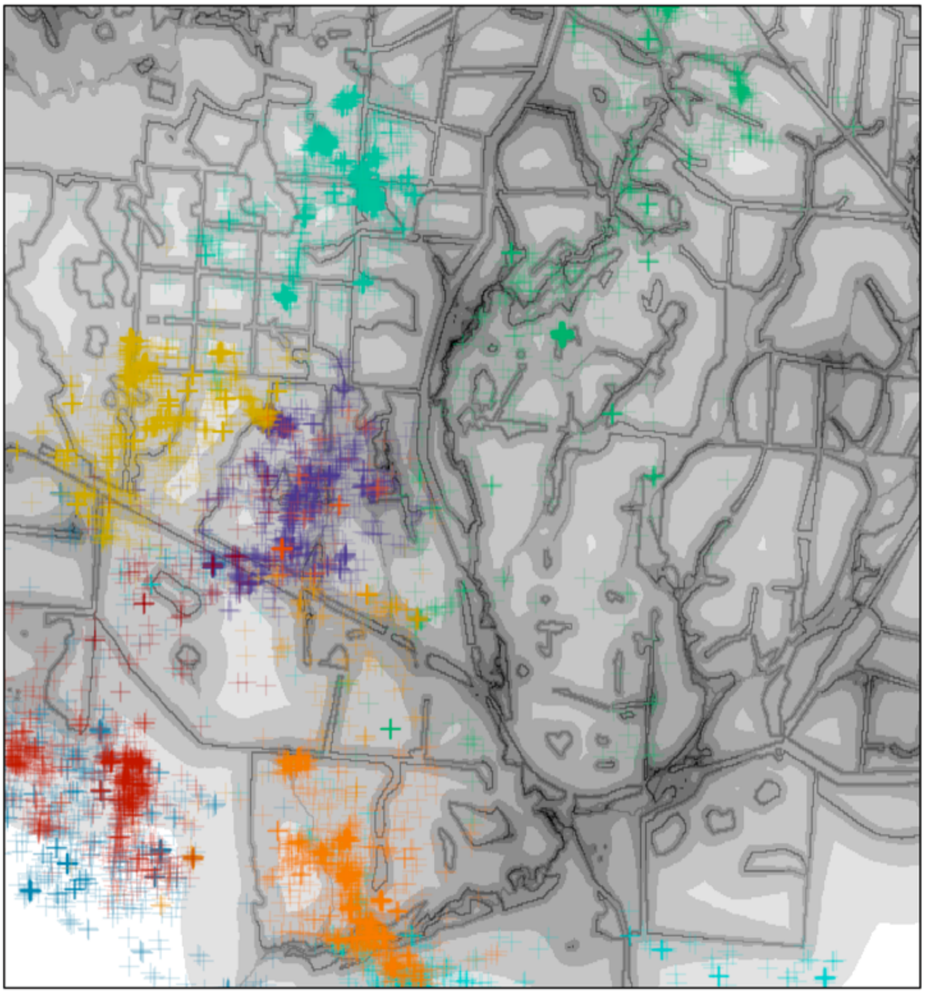

**Figure S1. Feral cat locations.** GPS-locations of 6 adult female (warm colours) and 5 adult male feral cats (cool colours). The background represents the resource map, which was derived by calculating the 1–($\log_2$+1) scaled Euclidean distance from three environmental layers (water courses, vegetation and buildings).

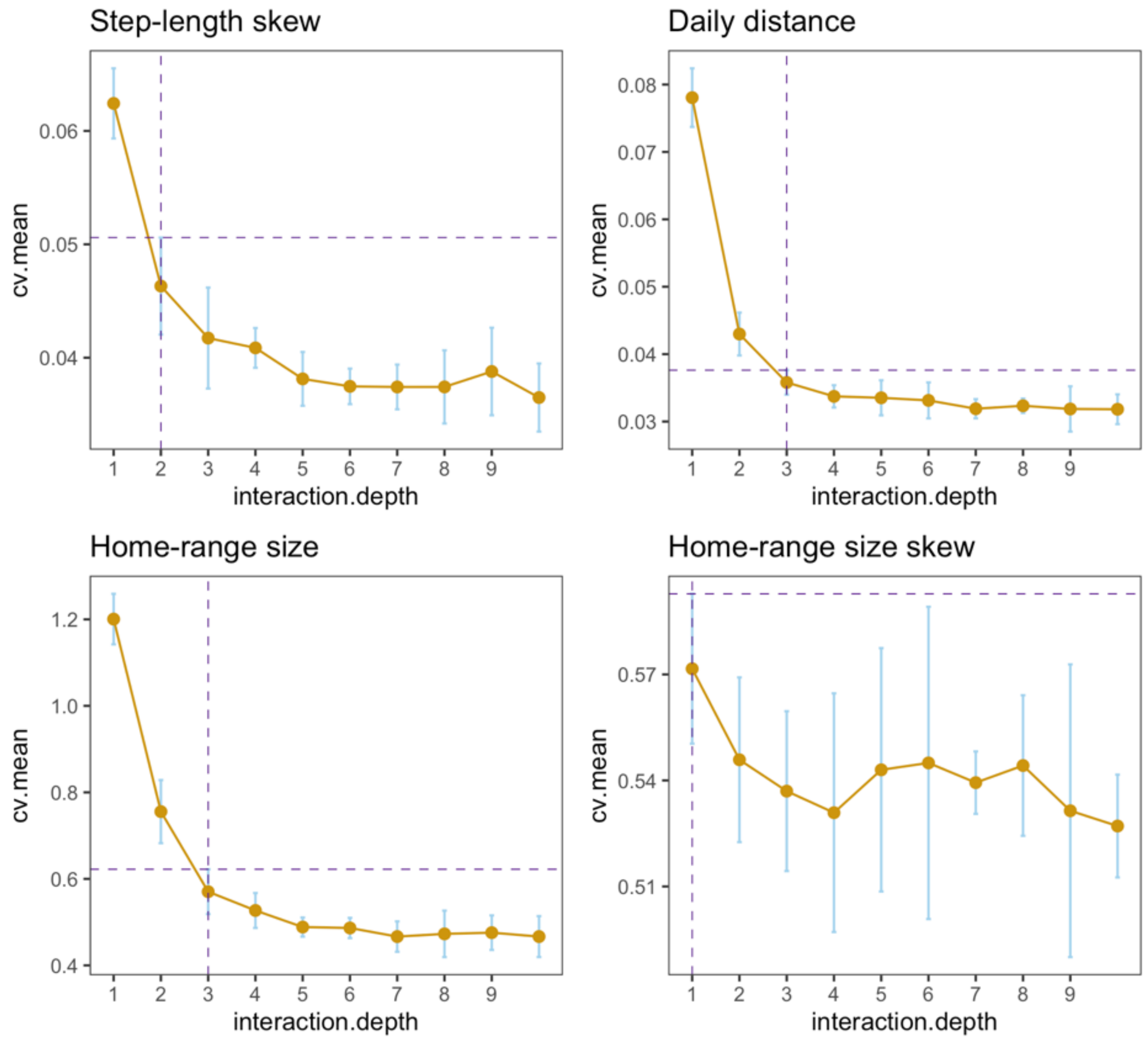


**Figure S2. Estimated prediction error curves and their standard errors for boosted regression tree (BRT) models of each summary statistic.** Each curve is plotted as a function of interaction depth (i.e., tree complexity). The estimates of prediction error and their standard errors were obtained by fivefold cross-validation ('CV'). The least complex model within one standard error of the best is chosen, indicated by the purple vertical dashed lines. Note that we did not perform regularisation for step-length and HR overlap as CV deviance was less than 0.01 at tree complexity of 1.

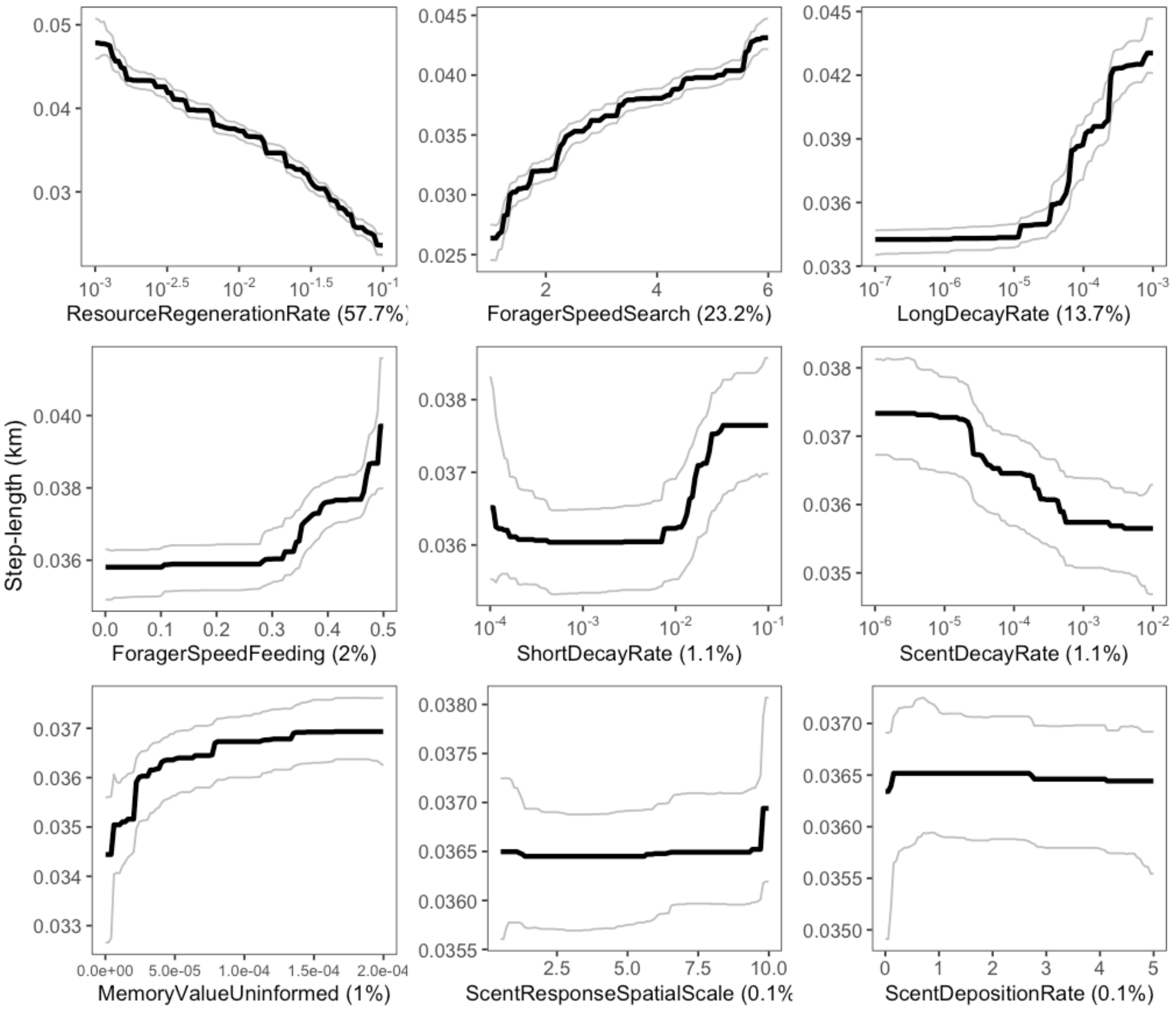


**Figure S3. Partial effects of input parameters on step-length** as generated from our boosted regression tree (BRT) analysis. Partial dependency plots show the effect of a given explanatory variable on step-length, while holding the effects of other explanatory variables at their average. We used 500 bootstrap replicates to calculate the 95% confidence intervals, depicted in grey. Note that several of the x-axes are on a log10 scale.

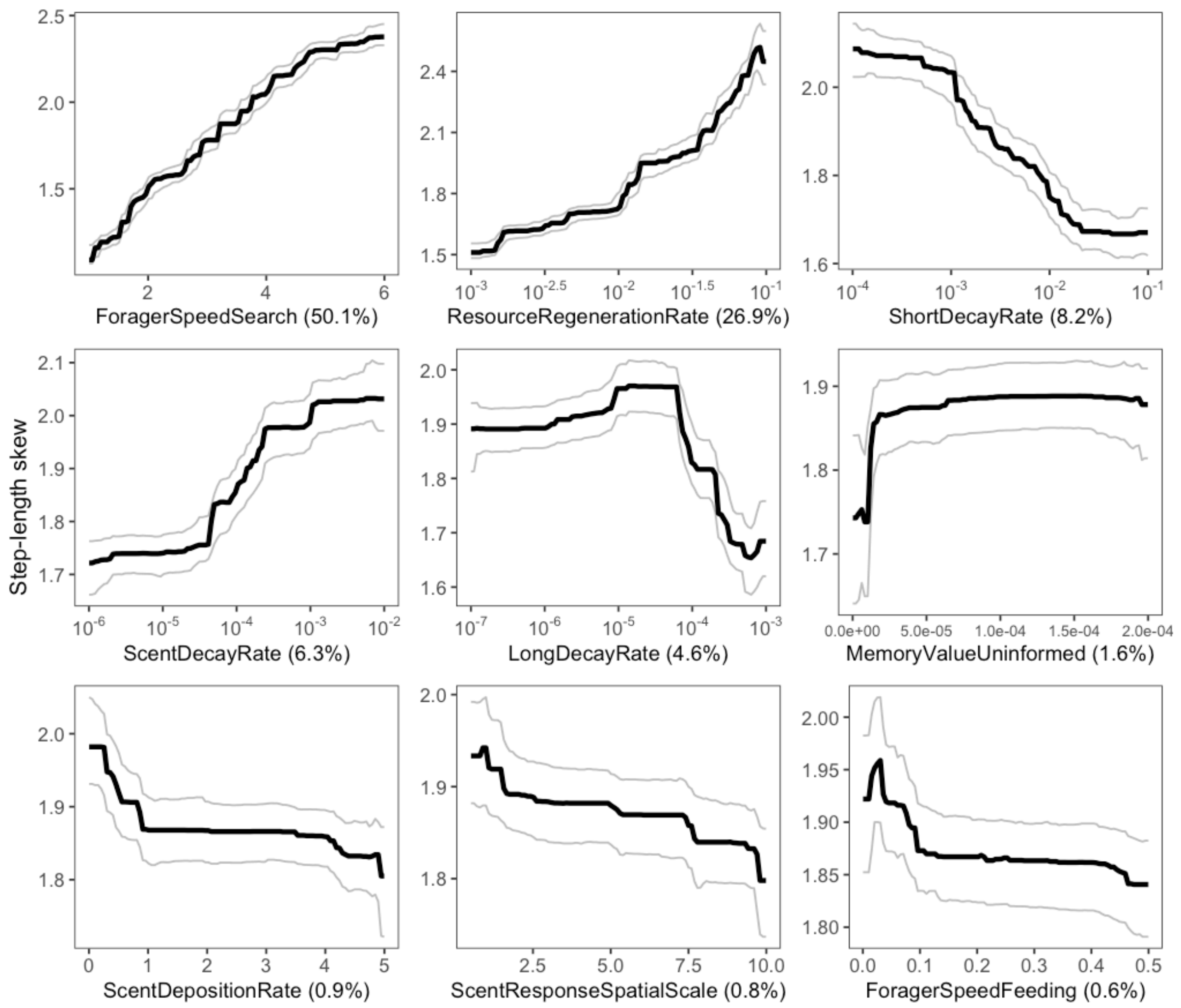


**Figure S4. Partial effects of input parameters on step-length skew** as generated from our boosted regression tree (BRT) analysis. Partial dependency plots show the effect of a given explanatory variable on step-length skew, while holding the effects of other explanatory variables at their average. We used 500 bootstrap replicates to calculate the 95% confidence intervals, depicted in grey. Note that several of the x-axes are on a log10 scale.

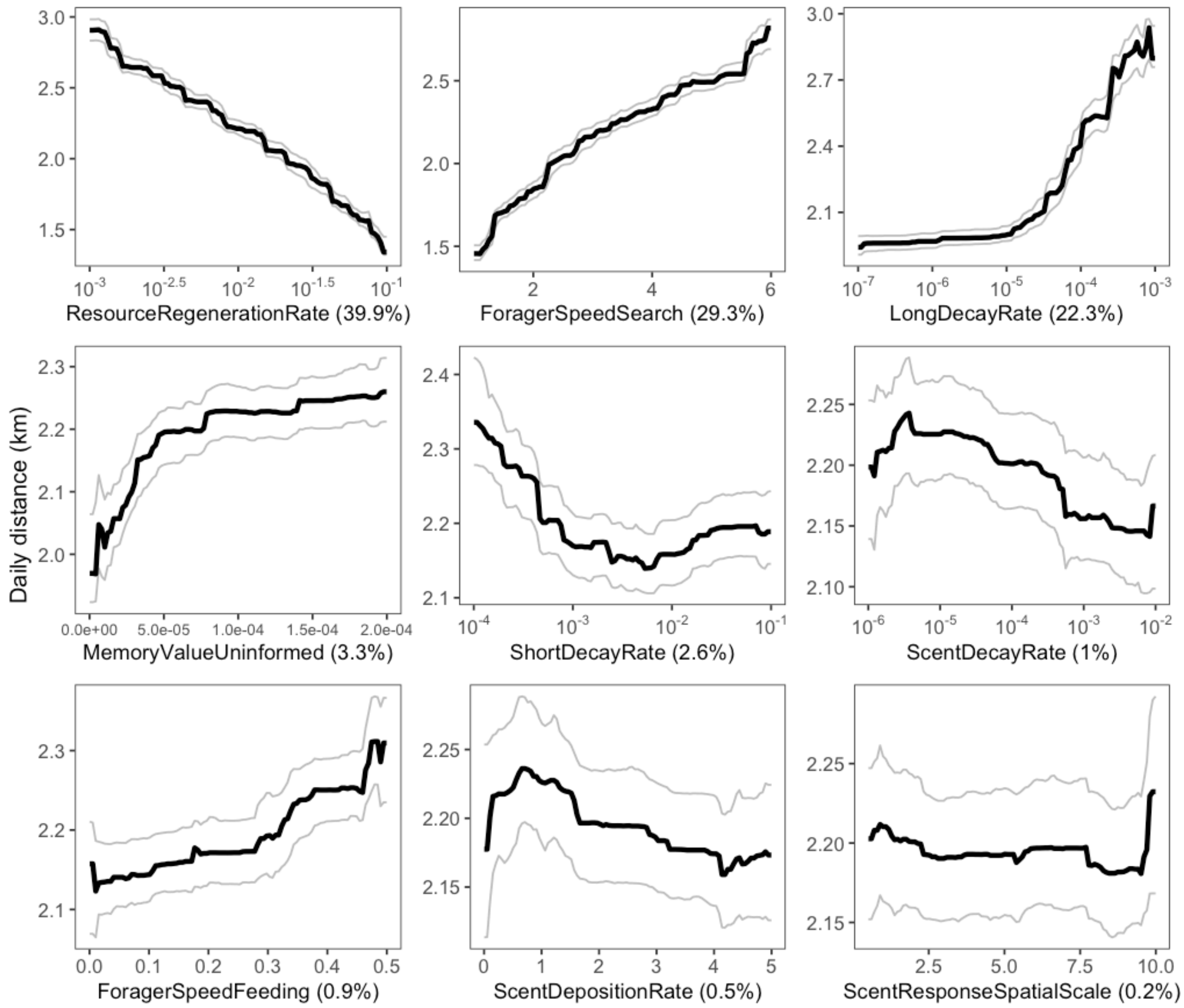


**Figure S5. Partial effects of input parameters on daily distance** (covered) as generated from our boosted regression tree (BRT) analysis. Partial dependency plots show the effect of a given explanatory variable on daily distance, while holding the effects of other explanatory variables at their average. We used 500 bootstrap replicates to calculate the 95% confidence intervals, depicted in grey. Note that several of the x-axes are on a log10 scale.

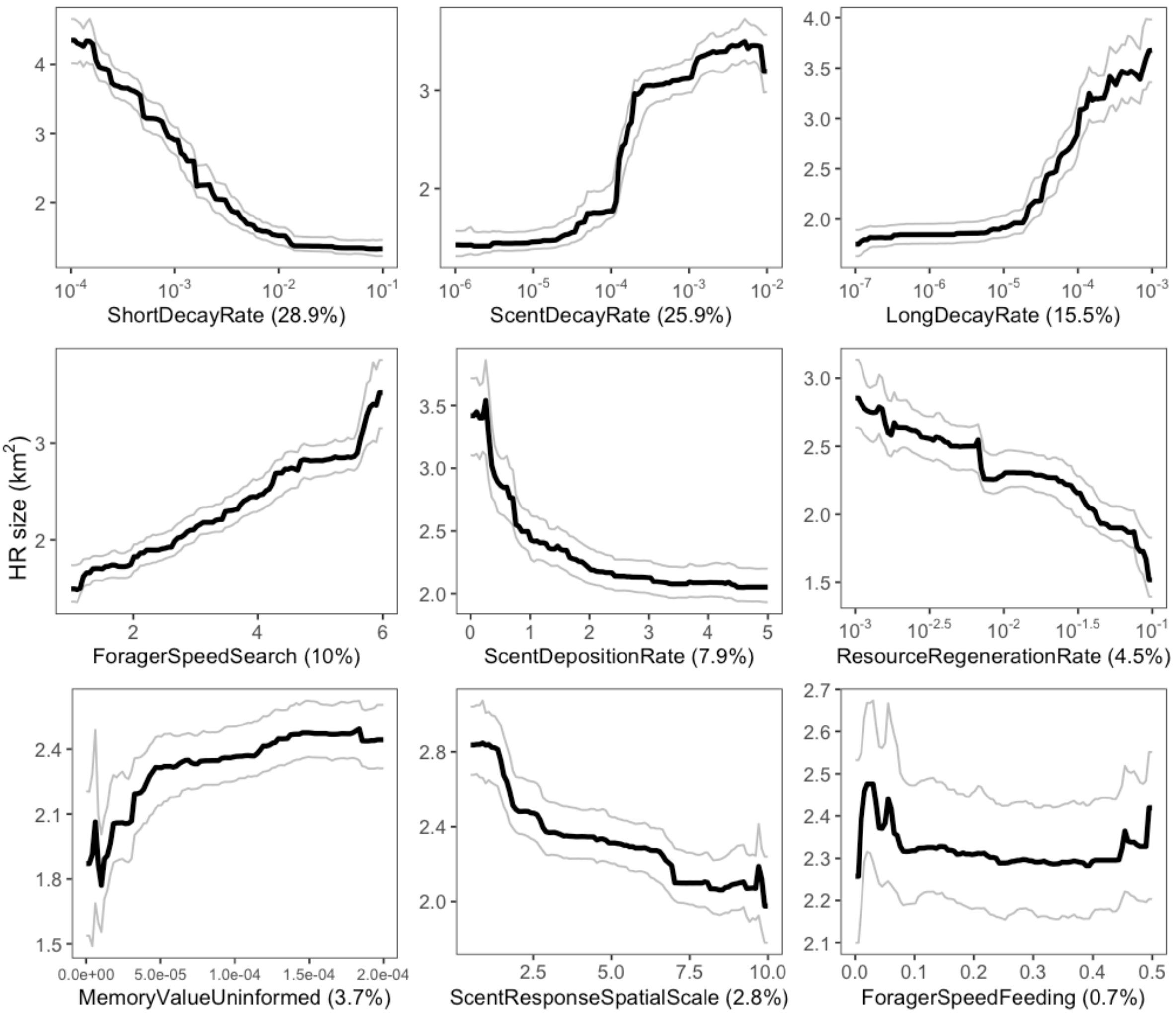


**Figure S6. Partial effects of input parameters on home-range (HR) size** as generated from our boosted regression tree (BRT) analysis. Partial dependency plots show the effect of a given explanatory variable on HR size, while holding the effects of other explanatory variables at their average. We used 500 bootstrap replicates to calculate the 95% confidence intervals, depicted in grey. Note that several of the x-axes are on a log10 scale.

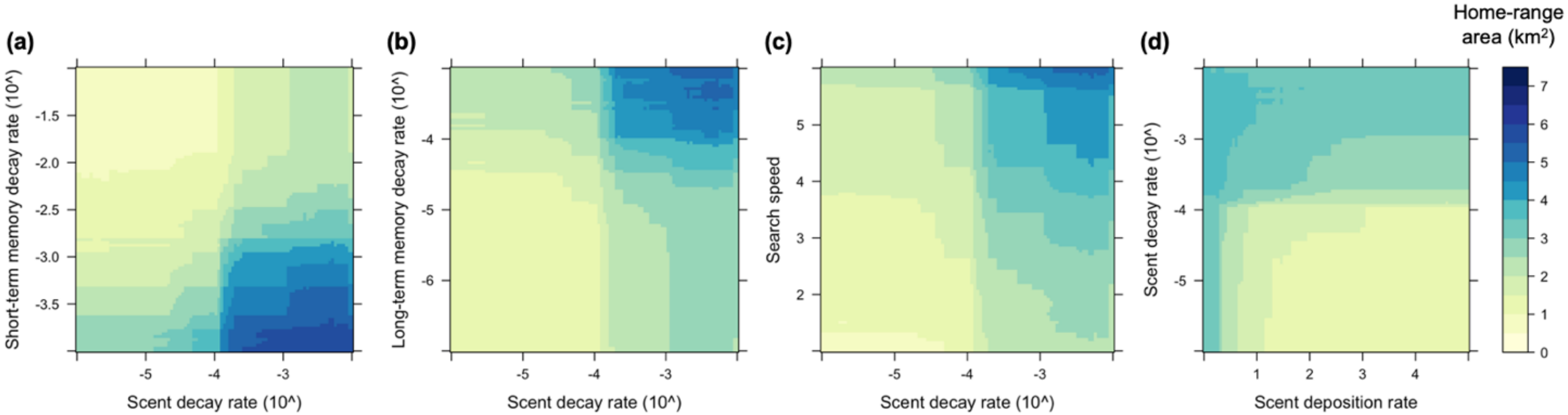


**Figure S7. Pairwise parameter interactions explaining home-range size,** including (a) scent decay rate and short-term memory decay rate, (b) scent decay rate and long-term memory decay rate, (c) search speed and scent decay rate, and (d) scent deposition rate and scent decay rate. Darker shades indicate larger home-range area as predicted from the boosted regression tree (BRT) model. Interaction sizes for each interaction are 75 (a), 43, (b), 34(c), 32(d).

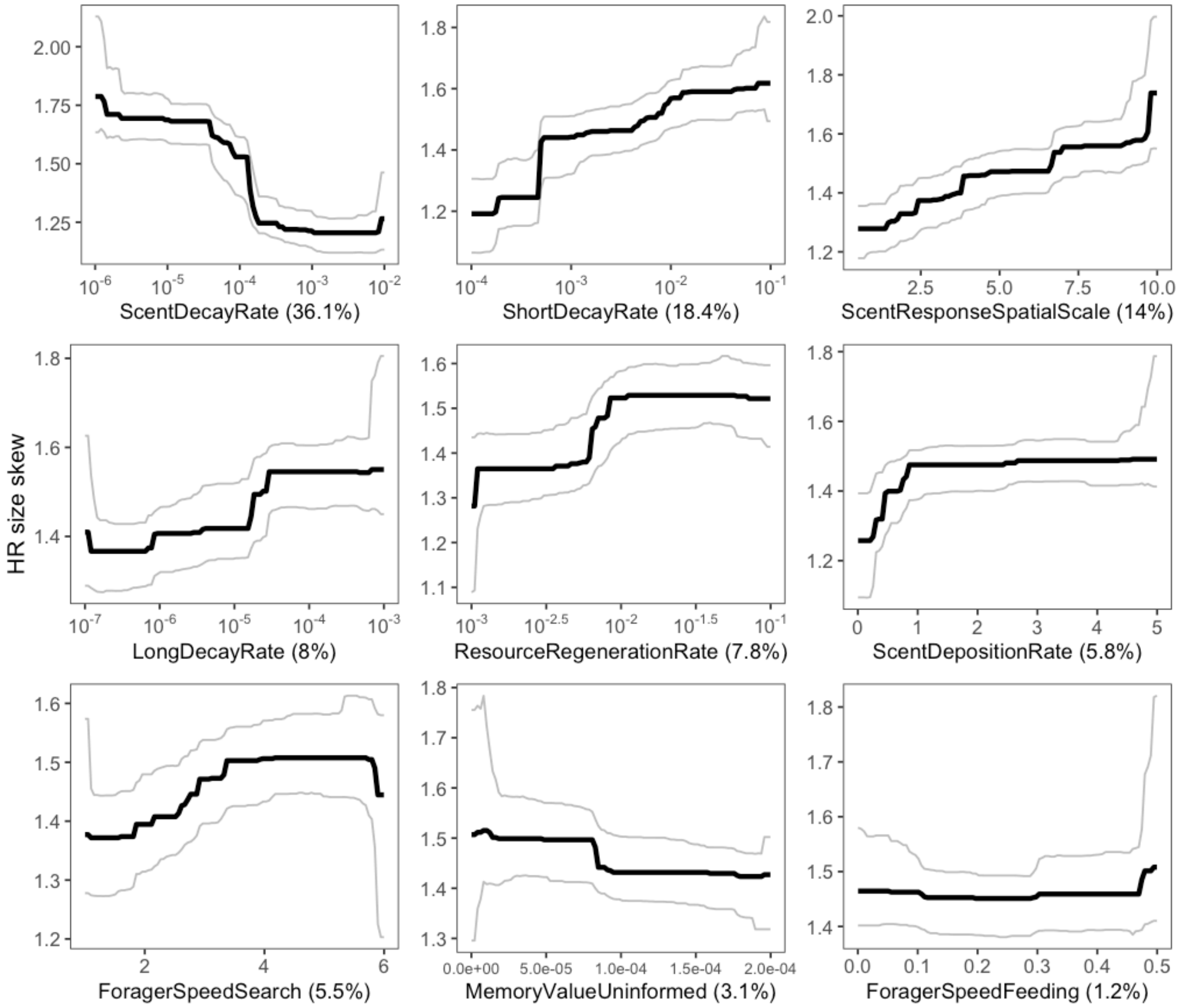


**Figure S8. Partial effects of input parameters on home-range (HR) size skew** (distribution skewness) as generated from our boosted regression tree (BRT) analysis. Partial dependency plots show the effect of a given explanatory variable on HR size skew, while holding the effects of other explanatory variables at their average. We used 500 bootstrap replicates to calculate the 95% confidence intervals, depicted in grey. Note that several of the x-axes are on a log10 scale.

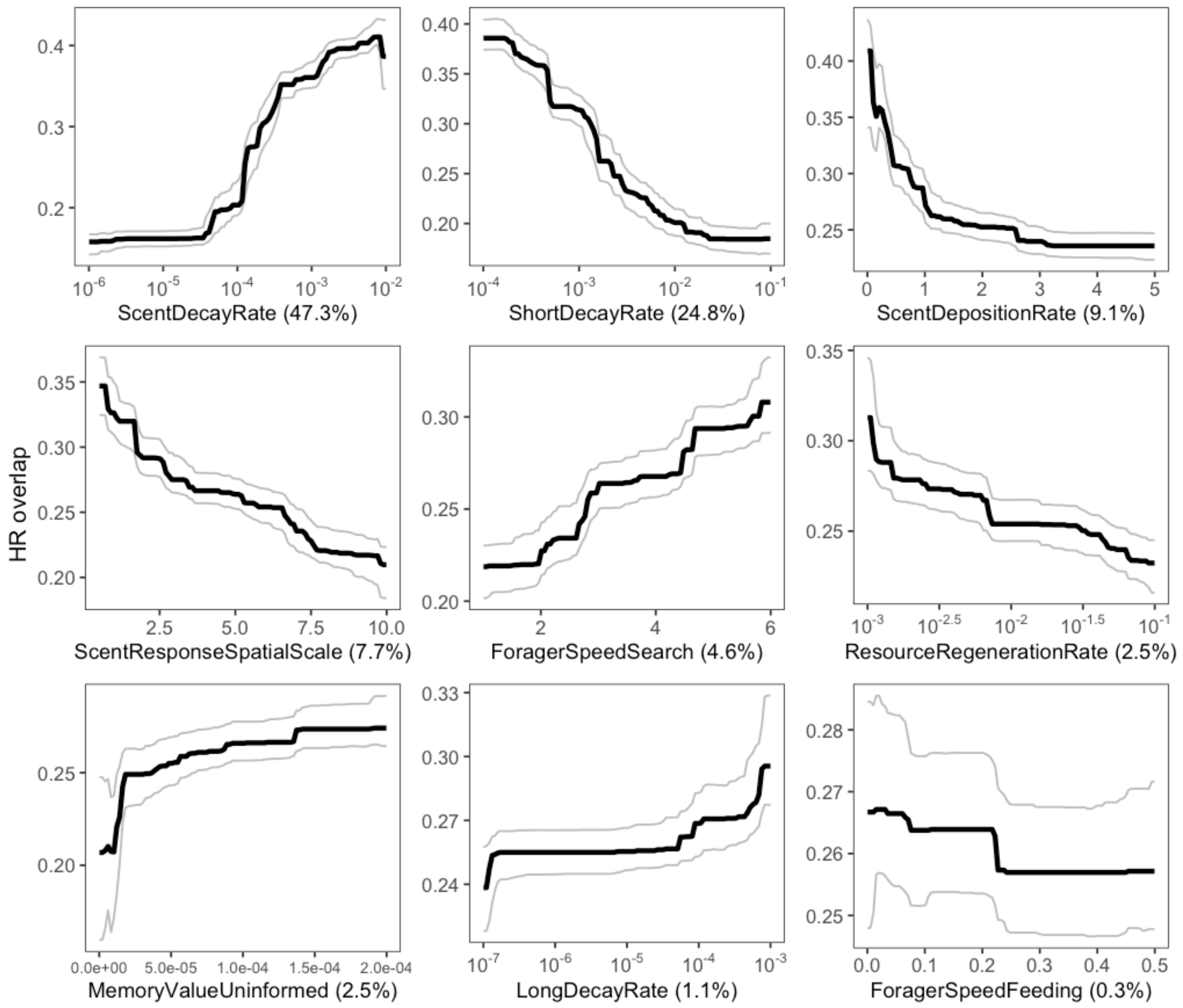


**Figure S9. Partial effects of input parameters on home-range (HR) overlap** (mean of maximum individual pairwise % overlap of 95% home-range area) as generated from our boosted regression tree (BRT) analysis. Partial dependency plots show the effect of a given explanatory variable on HR overlap, while holding the effects of other explanatory variables at their average. We used 500 bootstrap replicates to calculate the 95% confidence intervals, depicted in grey. Note that several of the x-axes are on a log10 scale.
